# Inferring antibiotic resistance selection in the environment can be confounded by correlations between resistance genes and unrelated functional traits

**DOI:** 10.1101/2025.10.12.681873

**Authors:** Cansu Uluseker, Sébastien Raguideau, Christopher Quince, Jan-Ulrich Kreft

## Abstract

Antimicrobial resistance (AMR) is a silent pandemic that is coupled with other crises such as climate change in the polycrisis humanity is facing. One of the key questions is whether antibiotic resistance genes (ARGs) are selected for at the low antibiotic concentrations typical for most environments. Many studies have observed changes in the relative abundance of ARGs from one environmental compartment to the next, e.g. from wastewater treatment plant influent to effluent. Fewer studies have directly tested for selection by incubating environmental samples in mesocosms or laboratory models at different concentrations of antibiotics to infer minimal selective concentrations. We developed a mathematical model to demonstrate that these studies can be confounded by shifts in the microbial community composition that occur when a microbiome is transported from one environmental compartment to another or when incubated under different conditions. Such community shifts will confound tests of selection when there is an association between carriage of ARGs and other functional traits. As an example, we show that there is a phylum-dependent association between the number of ARGs and the number of ribosomal RNA genes, which are both higher in fast growing, copiotrophic bacteria. We then show that specific growth rate or nutrient concentration upshifts increased the proportion of copiotrophs in the community and thus the relative abundance of ARGs. This result generalizes to community shifts for other reasons if there is some association between ARGs and ecological niches. Therefore, most studies of selection for ARGs in the environment are confounded. Solutions are proposed.

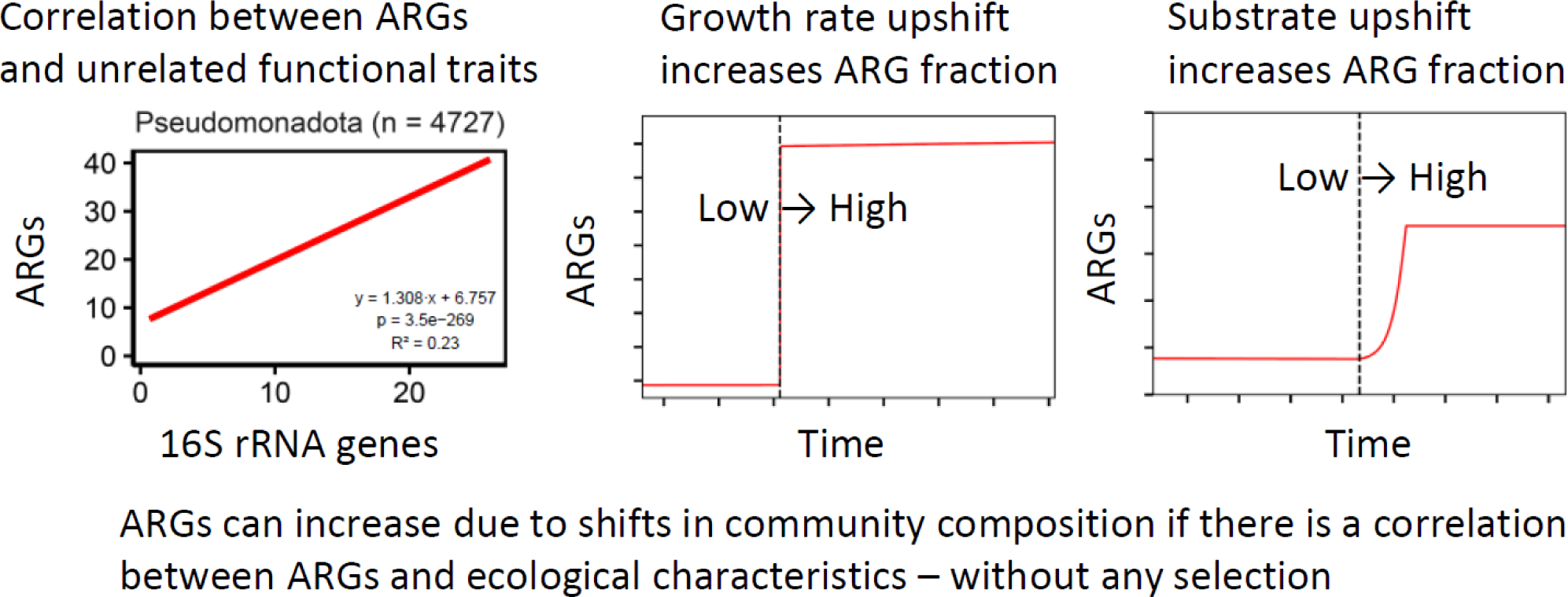

**Highlights:**

- ARG copy numbers are correlated with 16S rRNA gene copy numbers
- High rRNA gene numbers are typical for copiotrophs, which have high growth rates
- Hence, copiotrophs tend to have higher ARG numbers
- Changes in environmental conditions that increase copiotrophs increase ARGs
- Thus, ARGs can increase without selection due to shifts in community composition

## Introduction

Antimicrobial resistance (AMR) is a silent pandemic, less noticed than pandemics of rapidly spreading infectious diseases like COVID-19 and thus considered less urgent (Barron 2024). Like other pandemics, it has been spreading globally, causing a high death toll. The Global Burden of Disease (GBD) consortium study of 88 pathogen–drug combinations in 204 countries attributed 1·27 (95% UI 0·911–1·71) million deaths globally in 2019 to bacterial AMR (Murray *et al*. 2022). From their follow-up study of the period 1990-2021, they forecast an increase in mortality to 1·91 million (1·56–2·26) deaths attributable to AMR in 2050 (Naghavi *et al*. 2024). AMR, climate change, biodiversity loss and pollution are causally intertwined (United Nations Environment Programme (UNEP) 2023) and linked with other crises in the polycrisis humanity is currently facing (World Economic Forum (WEF) 2023, Lawrence *et al*. 2024).

The importance of including the environment in strategies to combat AMR has been recognized in the One Health concept (Larsson *et al*. 2023). The key environmental processes affecting the fate of resistant microbes are transport, growth and removal, selection and horizontal gene transfer (HGT) of resistance genes. Here, we focus on selection, which like the other processes is well understood in principle but not under the complexities of the environment. With the main exceptions of hospital and antibiotic manufacturing wastewater, concentrations of antibiotics in the environment are much lower than minimal inhibitory concentrations (MICs) and mostly lower than predicted no-effect concentrations for resistance selection (PNEC-Rs) derived from them (Bengtsson-Palme & Larsson 2016, Booth *et al*. 2020). Recent, more elaborate approaches to deriving PNEC-Rs or related measures (Emara *et al*. 2023, Kneis *et al*. 2025) produce wide distributions or confidence intervals that make these PNEC-Rs difficult to apply in practice. As a result, the question whether such low environmental concentrations of antibiotics are below the concentrations selecting for resistance under the conditions in the environment remains open.

One way to answer the question is to determine minimal selective concentrations (MSCs), which are concentrations where the growth rate of the resistant mutant (becoming higher than the sensitive wildtype at higher concentrations of the selective agent) intersects with the growth rate of the sensitive wildtype (higher in the absence of the selective agent due to a presumed fitness cost of resistance). Laboratory studies of resistant mutants competing with isogenic sensitive strains in rich media are highly sensitive and have shown that MSCs can be about 2 orders of magnitude lower than the corresponding MICs (Gullberg *et al*. 2011, 2014). In contrast, results from laboratory incubations of small environmental samples in rich media are highly variable (due to small sample volumes) and MSCs tend to be hard to discern and closer to the MICs (Murray *et al*. 2018, Stanton *et al*. 2020). More realistic studies using mesocosms, where large environmental samples are incubated, or lab-scale reactors that can mimic environmental conditions and maintain diversity to a larger degree, have found MSCs to be higher than in pure culture lab experiments (Muñoz-Aguayo *et al*. 2007, Knapp *et al*. 2008, Lundström *et al*. 2016, Tian *et al*. 2020). The higher diversity of the microbial communities in these systems has been shown to reduce selection for resistance (Klümper *et al*. 2019, Klümper *et al*. 2024, Fang *et al*. 2023).

Direct evidence for selection *in situ* is scarce and hard to obtain as it would require measuring resistance levels in the same microbial community over time (even if flowing away) as well as concentrations of potentially selective agents (which may fluctuate). Therefore, almost all studies in the environment only provide indications that selection may happen. For example, many studies have observed changes in the relative abundance of ARGs from one environmental compartment to the next, e.g. from wastewater treatment plant influent to effluent (Kim *et al*. 2010, Pallares-Vega *et al*. 2019, Honda *et al*. 2023, Park *et al*. 2024), and some also measured antibiotic concentrations (Gao *et al*. 2012, Mao *et al*. 2015). If the relative abundance of certain resistance genes increases, it is tempting to conclude that selection for resistance is responsible for this increase, even more so when concentrations of antibiotics were higher than in more pristine environments. For example, the relative abundance of fluoroquinolone resistant bacteria increased in a WWTP in Haridwar at fluoroquinolone concentrations of ∼10 μg/L (Kurasam *et al*. 2022).

However, we argue here that such evidence is likely to be confounded by shifts in the microbial community composition if there is an association between carriage of ARGs and ecological characteristics, regardless of the reasons for such an association. Community shifts inevitably occur when the community is transported from one compartment to another or when there is an inflow of another community. Community shifts are also impossible to avoid in laboratory studies where environmental samples containing a community of microbes are incubated under conditions that cannot fully reflect the environment the sample is taken from and may even be very far from environmental conditions, e.g. if a rich medium is added (Murray *et al*. 2018, Stanton *et al*. 2020). Community shifts are somewhat avoidable in mesocosms, especially if they are placed in the environment and large and complex enough to maintain the trophic structure and processes of the environment such as photosynthesis (Knapp *et al*. 2008).

Many studies have found changes in resistance levels and then erroneously taken an increase in resistance levels as evidence for selection, while others also looked for changes in community composition and some even recognizing the connection to shifts in resistance: Ma *et al*. (2024) found that an increase in filamentous bacteria, causing bulking in activated sludge, increased ARG levels because there are 1.5 times more ARGs per genome of filamentous bacteria compared to activated sludge bacteria overall. Guo *et al*. (2024) treated biofilms in tidal simulation mesocosms fed with water from the Yangtze estuary with silver nanoparticles for 28 days and found, using machine learning, structural equation modelling and metagenomic sequencing, that the nanoparticles reduced ARG levels by reducing betaproteobacteria as most ARGs were hosted by this taxon. Luo *et al*. (2022) found that addition of microplastics to lab-scale anaerobic digestors of activated sludge waste increased the relative abundance of ARGs because the microplastics enriched biofilm-forming bacteria, particularly *Acinetobacter* spp., which carry more ARGs. Forsberg *et al*. (2014) found that the soil resistome correlates with phylogenetic structure as ARGs were mostly nonmobile and as a result, nitrogen fertilisation shifted the community structure and ARG content. Zhao *et al*. (2018) used qPCR and 16S amplicon sequencing of faeces from pigs receiving different antibiotic feed additives in three large Chinese pig farms, finding that total ARG abundance correlated with antibiotic use, abundance of bacteria and mobile genetic elements (MGEs); they also found a correlation between specific ARGs and specific taxa, illustrating the complexities of inferring selection in the real world. Several other studies found that changes in community composition correlated with changes in the relative abundance of ARGs, e.g., when pig farm wastewater enters a river (Jia *et al*. 2017), between different compartments of WWTPs (Ju *et al*. 2019), or during thermophilic anaerobic digestion of microalgae-bacteria aggregates (Ovis-Sánchez *et al*. 2025). Variation in microbial community composition was the main reason for changes in ARG composition in an urban river polluted by point sources (Zhou *et al*. 2017). Zhu *et al*. (2025) found ARG composition in activated sludge from 142 WWTPs to correlate with particular taxa that are uncommon in the human gut. Similarly, diet changes causing shifts in community composition led to shifts in the gut resistome (Keskey *et al*. 2025). Often, there will be differences in community composition that are associated with differences in ARGs but drivers for those changes are unknown. Also, (partially) stochastic community assembly can produce spurious correlations in a particular habitat and at a particular time.

One common cause for community shifts when human or livestock wastes, full of AMR, enter the aquatic or terrestrial environment are downshifts in organic matter concentration. Such downshifts are expected to favour microbes adapted to resource-poor environments, known as oligotrophs, versus the microbes adapted to resource-rich environments, known as copiotrophs (Kuznetsov *et al*. 1979, Poindexter 1981, Koch 2001, Soler-Bistué *et al*. 2023). Oligotrophs are common in oligotrophic environments such as oceans (Schut *et al*. 1997, Button *et al*. 1998, Lauro *et al*. 2009) or bulk soils (Li *et al*. 2019, Dragone *et al*. 2024), though resource hotspots like plant roots can enrich for copiotrophs even in such environments. Copiotrophs will also be favoured when environmental samples are taken to the lab and supplemented with rich media. Roller *et al*. (2016) found that the copy number of the ribosomal RNA operon (*rrn*, consisting of 5S, 16S and 23S rDNA) in bacterial genomes increases with increasing maximal specific growth rate (log-log correlation), while the growth yield decreases. Thus, rrn copy number can be used to distinguish oligotrophic from copiotrophic bacteria, at least in a statistical sense. The copiotroph *E. coli* was found to be the main host of ARGs in wastewater, supporting the link between copiotrophs and resistance (Abdulkadir *et al*. 2024).

Here, we developed a simple mathematical model to demonstrate that observational and experimental studies of selection for resistance under environmental conditions can be confounded by shifts in the microbial community composition that inevitably occur when a microbiome is transported from one environmental compartment to another or when an environmental sample is incubated under different conditions. Since resource shifts are particularly relevant to AMR, the model was based on competing oligotrophic with copiotrophic bacteria. We hypothesized that copiotrophic bacteria such as *E. coli* are more likely to harbour ARGs and more competitive in high resource environments such that resource downshifts will lead to a shift from copiotrophic to oligotrophic bacteria, coupled with a decline in ARGs. Using complete genomes from 11,559 prokaryotic species, we show that there are indeed taxa-specific associations between ARG prevalence and *rrn* copy number, which in turn is associated with a copiotrophic growth strategy. Using our mathematical model, we then show that upshifts in specific growth rates or nutrient concentrations increased the proportion of copiotrophs in the community and thus the relative abundance of ARGs. This result generalizes to other reasons for community shifts as long as there is some association between ARGs and ecological niches. Therefore, most studies of selection for ARGs in the environment are confounded. We propose potential solutions for this problem.

## Materials & methods

### Bioinformatic analysis of ARG copy number association with rRNA gene copy number

The RefSeq database (Goldfarb *et al*. 2025) was chosen to obtain the widest high quality genome collection. Using the metadata file from 01/2025, only complete prokaryotic genomes were kept. Finally, genomes were chosen by using ETE (v3) (Huerta-Cepas *et al*. 2016) to obtain species names and randomly picking one unique entry per species, resulting in a total of 11,559 genomes. Krona (Ondov *et al*. 2011) was employed to visualize the taxonomic distribution of the selected 11,559 genomes.

All genomes were processed through gene calling using Prodigal (Hyatt *et al*. 2010) followed by AMR annotation using DIAMOND (v2.0.9) with option --more-sensitive (Buchfink *et al*. 2021) against the CARD database (v3.1.4) (McArthur *et al*. 2013). We used a best hits strategy but also filtered out any hits with (i) E-values higher than 1e-5, (ii) percentage identities smaller than 40% and (iii) alignment lengths smaller than 20% for either query or target. For this analysis, efflux ARGs were ignored.

Each genome was also annotated for ribosomal DNA using Barrnap (v0.9--3) (Seemann 2018). Whenever the number of identified 16S, 23S and 5S genes was not consistent, the average was taken.

### Model development and description

The ordinary differential equation (ODE) model was designed to simulate community shifts due to changes in environmental conditions while implementing an imperfect association between the ecological niches of the bacteria and their resistances.

Two environments were implemented in the model, a chemostat and a batch culture, as these simplified, unstructured environments can be viewed as the two extremes of a continuum along which most environments can be positioned, for a discussion of laboratory models of the environment see Wimpenny (1988). In fact, the layouts of gastrointestinal tracts of animals can be understood as different combinations of batch and chemostat reactors (Godon *et al*. 2013). Chemostats are open systems with a continuous inflow of resources and continuous removal of resources and biomass from the system, both at the same rate, the dilution rate (Herbert *et al*. 1956). The supply of resources into the system and removal of organisms capture fundamental characteristics of most environments and therefore chemostats and their variations have been used as models of a range of environments (Jannasch 1969, Wimpenny 1988, Finlay & Fenchel 1992, Jannasch & Egli 1993, Marsh 1995, Bull 2010). Due to negative feedback between growth and substrate concentration, chemostats reach a steady state where the specific growth rate equals the dilution rate (Herbert *et al*. 1956). Simply changing the dilution rate will change the system from an oligotrophic (low specific growth rate at low substrate concentration) to a copiotrophic environment (high specific growth rate at high substrate concentration). It is this shift that we are interested in here.

Batch cultures, on the other hand, are closed systems with an initial, often large, supply of resources and without any removal of resources and biomass, common in laboratories but also found in nature (e.g., dead organisms, fallen leaves, faeces). Here, we include batch cultures to represent those experimental studies of selection where environmental samples are taken to the laboratory to inoculate rich media under conditions far removed from the conditions of the sampled environment (Gullberg *et al*. 2011, Gullberg *et al*. 2014, Murray *et al*. 2018, Murray *et al*. 2020, Stanton *et al*. 2020).

Corresponding to the modelled chemostat at low dilution rates, we seeded our models with three oligotrophic bacteria, which were adapted to this low growth rate and low resource environment (e.g. clean river, ocean, forest soil) by having low maximal specific growth rates but high substrate affinities. Corresponding to the chemostat at high dilution rates, we also seeded our models with three copiotrophic bacteria, which were adapted to this high growth rate and high resource environment by having high maximal specific growth rates but low substrate affinities. These copiotrophs were also adapted to the batch culture. Oligotrophic bacteria were assumed to be less likely to carry resistance genes than copiotrophic bacteria, which we implemented by randomly assigning one out of three oligotrophic and two out of three copiotrophic bacteria to carry resistance genes. There was no antibiotic and thus no selection for resistance in these models.

The chemostat is a highly selective environment, with only one competitor for a shared, limited resource able to persist in the steady state, which is why chemostats have been used to understand this competitive exclusion principle (Armstrong & McGehee 1980). However, for our purposes, we needed to maintain our community and avoid competitive exclusion. This was achieved by providing six substitutable substrates – as many different resources as species. Additionally, the kinetics of the three oligotrophs were made distinct, yet similar, to ensure they were not identical yet remained oligotrophs. To this end, the maximum specific growth rates and substrate affinities for the different substrates were varied slightly. Likewise for the copiotrophs (**Table S1**).

Some species might become extinct in the chemostat; this will happen if their specific growth rate falls below the dilution rate at low concentrations of substrate (Tilman 1982). Once lost, a species cannot recover even if the dilution rate shifts up, so a constant low rate of immigration was included to continuously seed the system with each species.

We simulated two case studies. One where the chemostat environment changed from oligotrophic to copiotrophic by increasing the dilution rate ten-fold (mimicking a situation where bacteria are transported from one to another environmental compartment). This growth rate upshift will be referred to as **L2H** for “Low to High growth rate shift”. The other case study simulated a different shift from oligotrophic to copiotrophic, switching from chemostat to batch culture, by switching off the dilution rate and instead adding resources once at the same time. This growth rate upshift will be referred to as **E2L** for “Environment to Laboratory shift” as it corresponds to adding an environmental sample to rich medium in a batch culture.

We used the standard set of equations for chemostat dynamics (without maintenance) but included an immigration rate. For the specific growth rates, we used the modified Monod model for multiple substitutable substrates developed by Lendenmann & Egli (1998). All equations and parameters are provided in the **Supplementary text 1. Figure S2** visualizes the growth kinetics of the six species for the growth rate upshift case L2H while **Figure S5** visualizes the slightly different kinetics of species for the environment to laboratory shift case E2L.

## Results

### ARG numbers are associated with rRNA gene copy numbers in several phyla

Using the RefSeq database, we annotated AMR genes and ribosomal genes in complete genomes of 11,559 unique prokaryotic species. As a consequence of research bias, visualized using Krona (**Figure S1**), only 12 phyla contained more than 50 species with complete genomes, with Pseudomonadota, Actinomycetota and Bacillota containing the largest numbers of known genomes. The same phyla likely contain the largest fraction of copiotrophs.

While simply fitting the number of ARGs against the average number of ribosomal genes across all genomes did detect a significant correlation, a much better fit is obtained by assessing the correlation at the phylum level (**Figure 1**). Several phyla showed a clear and significant correlation, while others did not. It is likely that stronger correlations can be found for some phyla at a finer taxonomic level. For our argument to be valid, it is not necessary that all taxa show a correlation, only that changes in community composition have the potential to cause changes in relative ARG abundance. Indeed, if all taxa had the same correlation, the effects would cancel.

**Figure 1:**
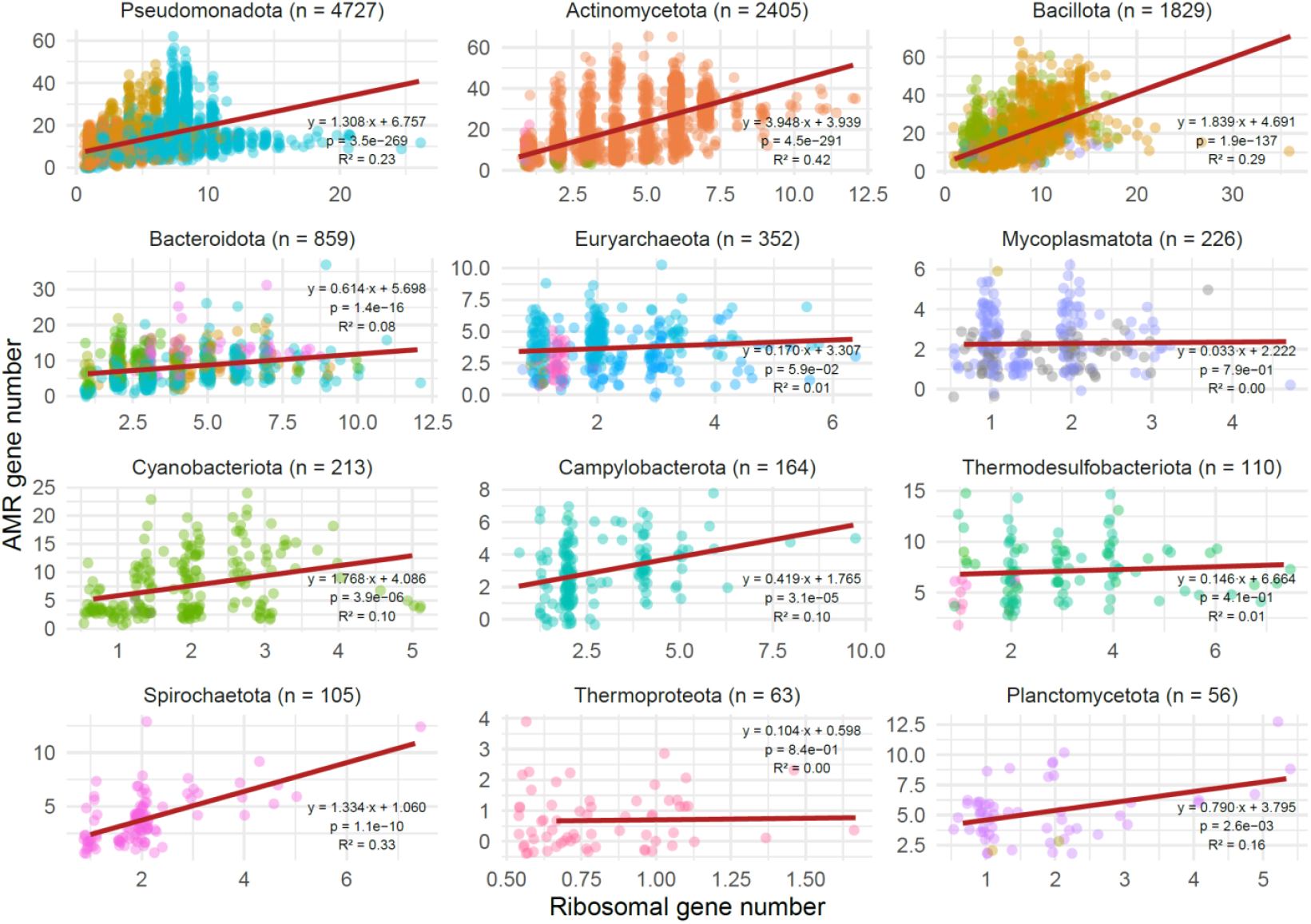
Ribosomal and AMR genes for several prokaryotic phyla showed a positive correlation. The correlation tends to be stronger and more significant for phyla with a wider range of ribosomal and AMR genes. Colour differentiates classes.

### Effects of growth rate upshifts

In this shift from low to high growth rate case study (L2H), the chemostat dynamics were simulated until the system had reached a steady state, which was characterised by low specific growth rates and substrate concentrations. Then, the dilution rate was increased, and a new steady state with higher specific growth rates (but only marginally higher substrate concentrations) was reached, as typical for chemostats operating well below the critical dilution rate (Herbert *et al*. 1956). In this new steady state, the copiotrophs were dominant and as a result of the link between growth rate and resistance, the fraction of resistant bacteria had increased (**Figure 2**). In this case, also the absolute abundance of resistant bacteria had increased (**Figure S3**). This is because in the chemostat, the population densities depended primarily on the concentrations of substrates in the inflow (if the concentration in the chemostat itself is negligible as in this case study, see **Figure S4**) times the (fixed) growth yields. Since the steady states remain the same (they are attractors) regardless of the direction or order of shifts, the reverse is also true, i.e., the resistance fraction would drop if the dilution and thus specific growth rate were decreased.

**Figure 2:**
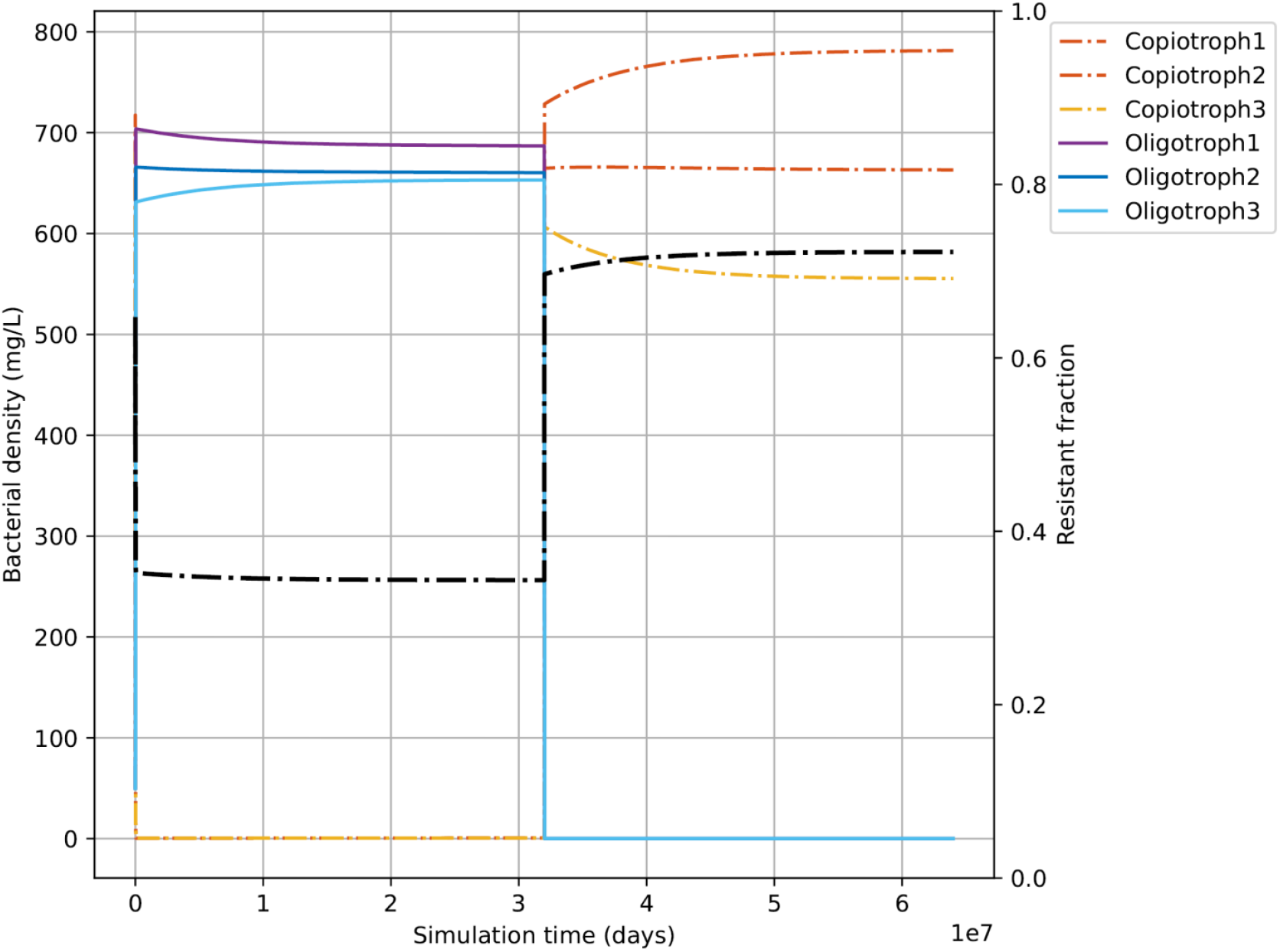
An upshift in the chemostat’s dilution rate led to a shift from oligotrophic species (blue colours) to copiotrophic species (red colours) and an increase in resistance. As two out of three copiotrophic species and only one out of three oligotrophic species carried a resistance gene, the relative abundance of the resistance gene (black line) increased following the upshift. In this case, also the absolute abundance of the resistant bacteria increased (**Figure S3**). The chemostat dynamics was simulated long enough to reach a steady state before and after the upshift, with low concentrations of the substrates before and after the upshift (**Figure S4**).

### Effects of substrate upshifts

The environment to laboratory case study (E2L) was modelled as a shift from a chemostat with continuously low growth rates and low substrate concentrations, capturing characteristics of most environments, to a batch reactor that supports fast growth for a short period of time. The first part of this was essentially the same as the first case study (but not identical as kinetic parameters were slightly adjusted). The chemostat was simulated until a steady state was reached where the specific growth rates matched the dilution rate of the system. Due to low specific growth rates and low substrate concentrations, the oligotrophs with their higher substrate affinity and lower specific growth rate potential were favoured and dominated the community (**Figure 3**). Copiotrophs were maintained in the chemostat only because of a baseline immigration rate.

**Figure 3:**
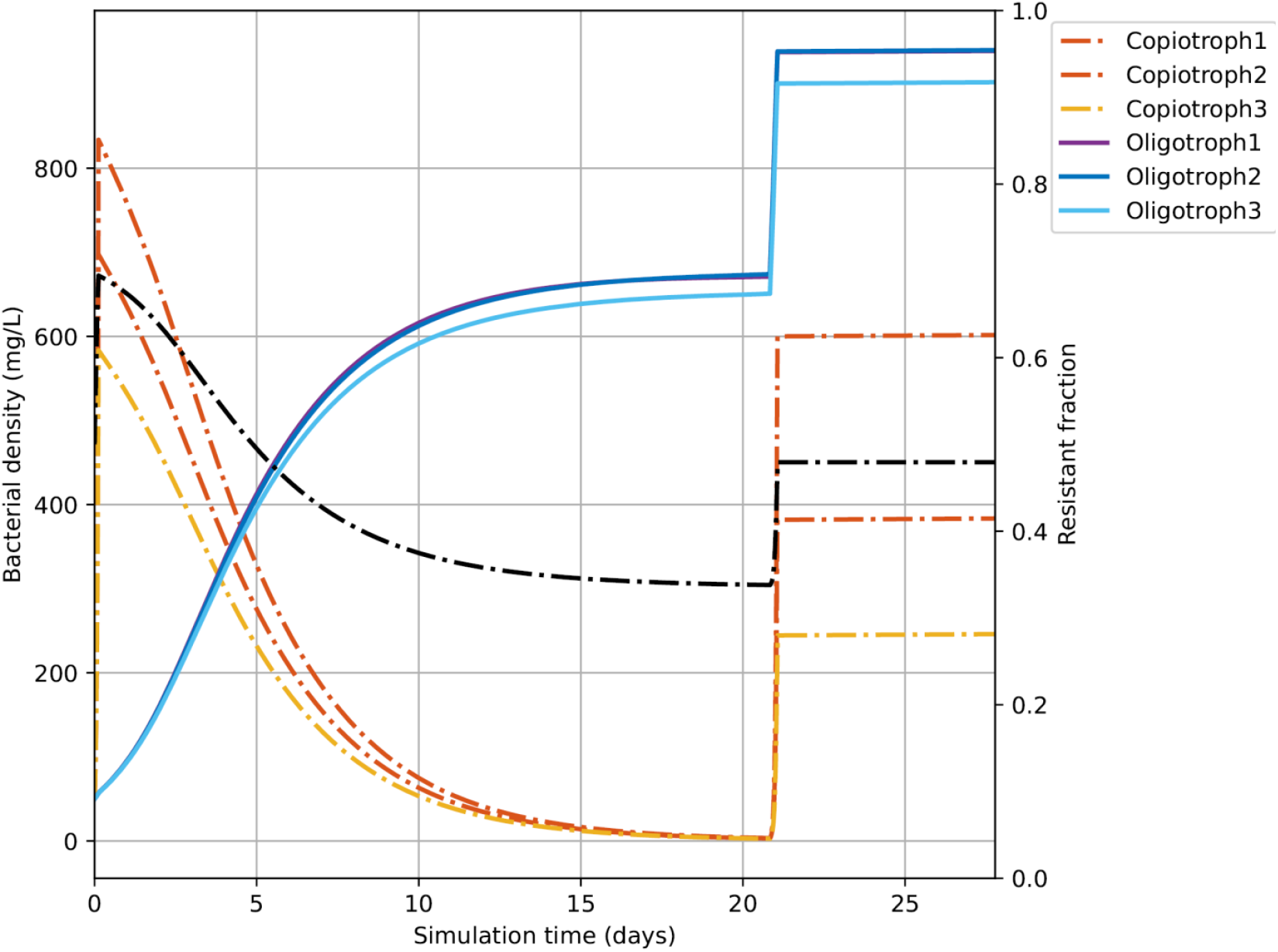
The fraction of resistant bacteria (black line) increased when simulating the E2L case study. Here the shift was from a chemostat system with low growth rates and substrate concentrations, typical for most environments, to a batch culture with initially high substrate concentrations, typical for some environments and laboratory models. Since bacteria were not removed in the batch phase, both oligotrophs and copiotrophs increased, albeit the latter more strongly, which caused an increase in the relative abundance of the resistance gene. Also in this case, the absolute abundance of the resistant bacteria increased (**Figure S6**). Since the population in the batch culture reached a higher density than in the chemostat, the increase in absolute abundance was above a proportional increase in relative abundance. The chemostat dynamics was simulated long enough to reach a steady state before the shift and then the batch culture was simulated until the substrates were consumed (**Figure S7**) and growth ceased.

Once the steady state had been reached, the continuous supply of nutrients and removal of biomass was turned off (the dilution rate was set to zero), and one shot of a large amount of all substrates was given (equivalent to inoculating a batch culture with a sample from the chemostat representing the environment). Despite the oligotrophs dominating the ‘inoculum’, the temporarily high substrate concentrations in the batch reactor enabled the copiotrophs to grow faster than the also growing oligotrophs until most of the substrate had been depleted. While oligotrophs had a higher specific growth rate in this last phase of batch growth, the small amount of substrate left then did not allow a large increase in biomass for any of the competitors. Therefore, the copiotrophs increased more than the oligotrophs (**Figure 3**), and as a result the absolute and relative abundance of resistant bacteria increased (**Figure S6**). Since our simple model does not contain any maintenance or death terms, a ‘frozen’ steady state was reached where populations remained constant, in contrast to the dynamic and stable steady state of the chemostat.

## Discussion

### Community shifts affect resistance levels

In both case studies, the switch favoured the copiotrophs and with their relative and absolute increase in the community, the relative and absolute abundance of ARGs increased. The reverse, switching from copiotrophic to oligotrophic environments, will lead to a drop in ARGs. Thus, community shifts can confound observational studies of selection in the environment where connected environmental compartments are compared, but also experimental studies where resource levels or other conditions change, resulting in community shifts. Mesocosms that better maintain environmental conditions should be the least confounded type of study.

Considering the flow of ARGs from human and animal origins through the environment, the typical situation is a drop in resource levels, which should result in a shift from copiotrophs to oligotrophs in these communities and therefore a drop in ARGs. Examples are the dilution of human faeces entering sewers, followed by further drops in resource levels when the sewage enters treatment plants and then rivers. Similarly for animal faeces entering slurry tanks or manure heaps and then again upon spreading on soils. Indeed, levels of ARGs decrease along those pathways (Lin *et al*. 2022, Martiny *et al*. 2022).

### Absolute versus relative abundance

We have focussed so far on relative abundance since almost all studies report changes in relative abundance of ARGs while not all report both relative and absolute abundances. If the total community size does not change from one compartment to the next, changes in relative and absolute abundance are equivalent. However, it is almost inescapable that the total community size changes so it is important to consider how this will affect relative and absolute abundances of ARGs. If total community size increases, absolute abundance obviously increases if the relative abundance increases, but it can also increase if the relative abundance decreases. Inversely, if total community size decreases, the absolute abundance obviously decreases if the relative abundance decreases, but it can also decrease if the relative abundance increases.

### Associations between ARGs and ecological niches are not limited to copiotrophs

We have used the association between ARG prevalence and copiotrophic growth strategy as an example, but any other association between resistance(s) and ecological niche(s) likely results in changes in the relative abundance of ARGs when the community composition changes. For example, shifts in community composition due to changes in oxygen and temperature along an urban river in the Yangtze watershed were coupled to decreasing ARG abundance (Wu *et al*. 2023). In soil mesocosms, Li *et al*. (2025) have shown that higher plant diversity, leading to higher resource diversity in root exudates, shifts the soil microbiome from bacterial hosts that harbour abundant ARGs and MGEs to those that do not, leading to a reduction in ARG and MGE abundance.

### Colonization resistance can alter resistance levels without resistance selection

Whether due to directed transport of microbial communities from one compartment to another, via wastewater pipeline or river or rainfall-runoff, or due to stochastic dispersal, individual microbes or whole communities enter resident communities all the time. If the resident community includes microbes that are ecologically similar, such as sensitive bacteria that are relatives of invading resistant bacteria, a priority effect, known in gut microbiology as colonization resistance, explains the hurdles faced by invaders (Freter *et al*. 1986, Brugiroux *et al*. 2016, Eberl *et al*. 2021, Letten *et al*. 2021, Segura Munoz *et al*. 2022). A study of immigration success of sewer communities from WWTP influents into lab-scale reactors inoculated with activated sludge found both replacement of the indigenous strains or failure of immigration, with the relative abundance of 11/15 ARGs in the activated sludge increasing (Gibson *et al*. 2023). If the resident community has a higher biodiversity, more invaders will be excluded, causing a barrier to invasion by resistant bacteria (Klümper *et al*. 2024).

### Community shifts can be exploited to reduce AMR

Community shifts often occur naturally, through transport of microbiomes or changes in environmental conditions in place, or they are engineered, such as in waste treatment processes, biostimulation or faecal microbiota transplantation (FMT). While the natural shifts often reduce AMR, engineered shifts could be specifically chosen or optimised to reduce AMR. When freshwater enters estuaries and then the marine environment, community shifts are inevitable and likely to result in changes in ARG levels. A global comparison of ARG prevalence across different biomes found lower prevalence in marine water compared to freshwater as would be expected from the many differences between marine and warm-blooded animal habitats (Lin *et al*. 2022). The microbial community residing in the colon experiences its first major shift upon defecation, when the environment changes from anaerobic to aerobic, as well as drops in temperature and downshifts in organic material. Later, entering wastewater treatment plants, further shifts can occur when the treatment process involves alternating anaerobic, anoxic and aerobic compartments and these shifts have been found to reduce AMR levels (Christgen *et al*. 2015). Treatment processes can also involve temperature shifts, e.g., from mesophilic microalgae-bacterial systems to thermophilic anaerobic digestion of sludge, leading to community shifts that result in a reduction of resistance (Ovis-Sánchez *et al*. 2025). However, some findings are inconsistent and are best compared by systematic reviews and metanalyses (Goulas *et al*. 2020). FMT partially replaces the gut microbiome of a recipient with that of a donor. If the donor is chosen to lack, e.g., multidrug-resistant bacteria, FMT can eliminate these bacteria through competition between closely related strains (Woodworth *et al*. 2023). As a corollary, ecological boundaries, across which shifts in community composition are expected to occur, have indeed been shown to affect the distribution of ARGs (Lin *et al*. 2022).

### Confounding due to upstream selection

Another pitfall that can lead to wrongly concluding that resistance has been selected in a particular environment is that the selection may have happened upstream of the environment studied. For example, the study by Brown *et al*. (2024) compiled data on community antibiotic prescription frequencies and combined this with an extensive timeseries of deep metagenomes in the influent of a WWTP resulting from COVID surveillance, to find that ARGs hosted by Enterobacteriaceae are associated with prescription frequency. However, selection of these ARGs by antibiotics could have happened upstream, in the guts of patients, rather than the sewers. This is in fact more likely as antibiotic concentrations in the patients are higher than in sewers. ARGs hosted by sewer indigenous Pseudomonadaceae were associated with prescription frequency after a delay of 1-3 months, which suggests more indirect effects than selection or possibly spurious correlations resulting from seasonal changes in prescription and seasonal changes in community composition that have independent causes and are out of sync. Moreover, they also found negative correlations between prescription frequency and relative abundance of ARGs, which could be an artefact of using relative abundance of ARGs as an absolute increase can become a relative decline when the community size increases while the composition shifts (Props *et al*. 2017, Vandeputte *et al*. 2017, Barlow *et al*. 2020, Ott *et al*. 2021). A more direct investigation of selection and HGT in lab-scale sequencing batch reactors fed with wastewater by the same lab did not find a consistent pattern of selection for resistance genes as most contigs with ARGs disappeared while some ARGs persisted in some contigs but not others, suggesting that their fate was coupled to the contig rather than selection (Brown, Maile-Moskowitz, *et al*. 2024).

### How can confounding be avoided?

We have argued that community shifts and upstream selection can masquerade as selection in the focal environment. But how can selection be evidenced directly? The most direct evidence would come from measuring changes in populations of isogenic strains where one strain is resistant while the other is sensitive, ideally the resistance gene would be the only difference between the strains, as in the seminal work by Gullberg *et al*. (2011). As a consequence, the strains would have nearly the same ecological niche, including competitors, predators, phages etc. This approach has been successfully used, for example by Klümper *et al*. (2019), finding that the presence of a pig faecal community increased the MSC of *E. coli* due to increased fitness cost of the resistance and community protection of the sensitive phenotype. Their following study showed that this ‘community effect’ is surprisingly robust against perturbation of the community by another antibiotic (Fang *et al*. 2023). In a synthetic community of isogenic sensitive and resistant *E. coli* with *Bacillus subtilis*, resistance selection was slowed down not due to competition for the single carbon and energy source glucose but due to the production of extracellular compounds by *B. subtilis* (Nair & Andersson 2023).

This approach has limitations since introducing such an isogenic pair into the environment would only be acceptable if at least one of them had been isolated from the environment and the other one had spontaneously arisen from resistance gene or plasmid loss or by de novo resistance mutation rather than by genetic engineering (though a clean deletion could be considered). However, such isogenic pairs may already exist in the environment studied. It is also possible to incubate strains in the environment in some kind of ‘cage’ that combines physical isolation with exposure to, and metabolite exchange with, the environment. For example, Lessard & Sieburth (1983) exposed *E. coli* and enterococci populations from raw sewage in a stirred, transparent cage immersed in an estuary and salt marsh to measure ‘decay’ rates. Natural or engineered isogenic strains could be used in mesocosms that by and large maintain the composition and diversity of an environmental community and reflect the relevant physicochemical conditions. Such mesocosm studies allow experimental manipulation of the concentrations of selective or co-selective compounds but have usually not been used with isogenic pairs (Knapp *et al*. 2008, Quinlan *et al*. 2011, Baker *et al*. 2022).

In the absence of isogenic pairs of resistant and sensitive bacteria, there is another avenue to evidence selection for resistance, which is also based on an isogenic background. Quintela-Baluja *et al*. (2021) designed qPCR primers to distinguish ARG carrying, from empty, clinical class 1 integrons, finding that these integrons carried 10 times more ARGs in hospital wastewater than in other compartments. Integrons in recycled activated sludge and the receiving river had lost ARGs. Since the only difference was carriage of ARGs, this is direct and clear evidence for selection for ARGs in hospital wastewater or ‘upstream’ in patients, and lack of selection in activated sludge and receiving rivers. Similarly, selection could be tested by tracking the fate of ARGs embedded in their (meta)genomic contexts while these contexts are transported from one environmental compartment to another, such as from gut microbiomes under antibiotic treatment to gut microbiomes not under treatment (Lee *et al*. 2023).

We are not aware of alternative avenues to directly test for resistance selection. Perhaps, indirect evidence that is at risk of confounding could be used if sufficiently large studies encompassing different and known levels of potentially confounding factors enable statistical analysis that can control for these factors.

## Conclusions and recommendations

Our finding that microbial community shifts can result in changes of absolute and relative abundance of ARGs and ARBs in the absence of selection (and HGT) is trivial in the sense that it follows logically from the assumed coupling of resistance with ecology. This coupling is well documented though complex and variable in its details.

Our point is that the consequences of this trivial result are far from trivial, as any observational or experimental study of changes in ARG abundance cannot provide unequivocal evidence for or against selection whenever there has also been a change in environmental conditions, which is hard to avoid. Any incubation of a microbiome sampled from the environment in the laboratory will undoubtedly change environmental conditions. Using larger mesocosms, especially when placed *in situ*, will reduce this shift and maintain more of the diversity of the source environment, but some change in conditions is unavoidable even in mesocosms.

While our examples were based on an association between ARGs and copiotrophs (fast-growing bacteria), coupled to shifts from eutrophic conditions favouring copiotrophs to oligotrophic conditions favouring oligotrophs, the conclusions are general. Any association between ARGs and bacteria with particular ecological niches can lead to changes in relative ARG abundance when environmental conditions change such that the community composition changes (which is almost inevitable).

An association between ARGs and ecological niches may often be linked to an association between phylogeny and ecology that has been observed on various levels of hierarchy. For example, Pseudomonadota (Proteobacteria) phylum members tend to be fast-growing, copiotrophic bacteria that also tend to harbour various plasmids including those with ARGs.

HGT of ARGs has the potential to break associations between ARGs and ecology, however, associations between phylogeny and resistance plasmids on the one hand and phylogeny and growth strategy on the other, mean that HGT within such taxa may not break the association.

The issue of selection being confounded by community shifts prompts the question of how to obtain direct, unequivocal evidence of selection for or against resistance at certain concentrations of antibiotics or other selective agents?

Crucial for any solution is to somehow eliminate ecological differences between sensitive and resistant bacteria. Ideally, one would determine changes in relative abundance of a sensitive bacterium competing with a resistant bacterium that is otherwise isogenic, such as a resistant mutant of the same strain. This ought to guarantee that their ecology is very similar if not identical, though the presence of a resistance gene or plasmid has the potential to perturb gene expression and changes in ecological niche upon acquiring resistance have been reported (Letten *et al*. 2021). Spiking such an isogenic pair into laboratory or mesocosm incubations and tracking their abundance would then enable direct measurement of selection. Keeping track of an isogenic pair in the environment is possible but will be harder. For environmental observations there is fortunately an alternative, based on detecting the gain or loss of resistance genes from mobile genetic elements in the same strain or genomic context when the strain or genome is transported from one environmental compartment to another, for an example see Quintela-Baluja *et al*. (2021). A tendency for a resistance gene to get lost from the mobile genetic element, strain or genome in the downstream compartment would indicate lack of selection for this resistance gene in the downstream compartment. Both options rely on the use of an isogenic background to reduce confounding through ecological differences.

## Author Contributions

Cansu Uluseker developed the model and code, simulated the model and made the corresponding figures: Formal analysis, Investigation, Methodology, Software, Validation, Visualization, Writing – original draft. Sébastien Raguideau did the bioinformatic and statistical analysis and made the corresponding figures: Data curation, Formal analysis, Investigation, Methodology, Validation, Visualization, Writing – original draft. Christopher Quince guided the bioinformatic and statistical analysis: Conceptualization, Methodology, Supervision. Jan-Ulrich Kreft came up with the idea for this study and guided the modelling and drafting of the paper: Conceptualization, Funding acquisition, Project administration, Supervision, Writing – original draft, Writing – review & editing.

## References

Abdulkadir N, Saraiva JP, Zhang J, Stolte S, Gillor O, Harms H, Rocha U (2024). Genome-centric analyses of 165 metagenomes show that mobile genetic elements are crucial for the transmission of antimicrobial resistance genes to pathogens in activated sludge and wastewater. Microbiology Spectrum 0: e02918–23 10.1128/spectrum.02918-23

Armstrong RA, McGehee R (1980). Competitive exclusion. American Naturalist 115: 151–170 10.1086/283553

Baker M, Williams AD, Hooton SPT, Helliwell R, King E, Dodsworth T, Baena-Nogueras RM, Warry A, Ortori CA, Todman H, Gray-Hammerton CJ, Pritchard ACW, Iles E, Cook R, Emes RD, Jones MA, Kypraios T, West H, Barrett DA, Ramsden SJ, Gomes RL, Hudson C, Millard AD, Raman S, Morris C, Dodd CER, Kreft J-U, Hobman JL, Stekel DJ (2022). Antimicrobial resistance in dairy slurry tanks: a critical point for measurement and control. Environment International 169: 107516 10.1016/j.envint.2022.107516

Barlow JT, Bogatyrev SR, Ismagilov RF (2020). A quantitative sequencing framework for absolute abundance measurements of mucosal and lumenal microbial communities. Nature Communications 11: 2590 10.1038/s41467-020-16224-6

Barron M (2024). The Antimicrobial Resistance Pandemic: Breaking the Silence. Microcosm

Bengtsson-Palme J, Larsson DGJ (2016). Concentrations of antibiotics predicted to select for resistant bacteria: Proposed limits for environmental regulation. Environment International 86: 140–149 10.1016/j.envint.2015.10.015

Booth A, Aga DS, Wester AL (2020). Retrospective analysis of the global antibiotic residues that exceed the predicted no effect concentration for antimicrobial resistance in various environmental matrices. Environment International 141: 105796 10.1016/j.envint.2020.105796

Brown CL, Maile-Moskowitz A, Lopatkin AJ, Xia K, Logan LK, Davis BC, Zhang L, Vikesland PJ, Pruden A (2024). Selection and horizontal gene transfer underlie microdiversity-level heterogeneity in resistance gene fate during wastewater treatment. Nature Communications 15: 5412 10.1038/s41467-024-49742-8

Brown CL, Rumi MA, McDaniel L, Maile-Moskowitz A, Sein J, Nguyen L, Choi M, Hindi F, Mullet J, Emon M, Moumi NA, Blair MF, Davis BC, Rao J, Baffoe-Bonnie A, Vikesland P, Pruden A, Zhang L (2024). Metagenomics disentangles epidemiological and microbial ecological associations between community antibiotic use and antibiotic resistance indicators measured in sewage. 2024.12.11.24318846 10.1101/2024.12.11.24318846

Brugiroux S, Beutler M, Pfann C, Garzetti D, Ruscheweyh H-J, Ring D, Diehl M, Herp S, Lötscher Y, Hussain S, Bunk B, Pukall R, Huson DH, Münch PC, McHardy AC, McCoy KD, Macpherson AJ, Loy A, Clavel T, Berry D, Stecher B (2016). Genome-guided design of a defined mouse microbiota that confers colonization resistance against Salmonella enterica serovar Typhimurium. Nature Microbiology 2: 16215 10.1038/nmicrobiol.2016.215

Buchfink B, Reuter K, Drost H-G (2021). Sensitive protein alignments at tree-of-life scale using DIAMOND. Nature Methods 18: 366–368 10.1038/s41592-021-01101-x

Bull AT (2010). The renaissance of continuous culture in the post-genomics age. Journal of Industrial Microbiology & Biotechnology 37: 993–1021

Button DK, Robertson BR, Lepp PW, Schmidt TM (1998). A small, dilute-cytoplasm, high-affinity, novel bacterium isolated by extinction culture and having kinetic constants compatible with growth at ambient concentrations of dissolved nutrients in seawater. Applied and Environmental Microbiology 64: 4467–4476

Christgen B, Yang Y, Ahammad SZ, Li B, Rodriquez DC, Zhang T, Graham DW (2015). Metagenomics Shows That Low-Energy Anaerobic−Aerobic Treatment Reactors Reduce Antibiotic Resistance Gene Levels from Domestic Wastewater. Environmental Science & Technology 49: 2577–2584 10.1021/es505521w

Dragone NB, Hoffert M, Strickland MS, Fierer N (2024). Taxonomic and genomic attributes of oligotrophic soil bacteria. ISME Communications 4: ycae081 10.1093/ismeco/ycae081

Eberl C, Weiss AS, Jochum LM, Raj ACD, Ring D, Hussain S, Herp S, Meng C, Kleigrewe K, Gigl M, Basic M, Stecher B (2021). E. coli enhance colonization resistance against Salmonella Typhimurium by competing for galactitol, a context-dependent limiting carbon source. Cell Host & Microbe 29: 1–13 10.1016/j.chom.2021.09.004

Emara Y, Jolliet O, Finkbeiner M, Heß S, Kosnik M, Siegert M-W, Fantke P (2023). Comparative selective pressure potential of antibiotics in the environment. Environmental Pollution 318: 120873 10.1016/j.envpol.2022.120873

Fang P, Elena AX, Kunath MA, Berendonk TU, Klümper U (2023). Reduced selection for antibiotic resistance in community context is maintained despite pressure by additional antibiotics. ISME Communications 3: 52 10.1038/s43705-023-00262-4

Finlay BJ, Fenchel T (1992). An anaerobic ciliate as a natural chemostat for the growth of endosymbiotic methanogens. European Journal of Protistology 28: 127–137 10.1016/S0932-4739(11)80041-6

Forsberg KJ, Patel S, Gibson MK, Lauber CL, Knight R, Fierer N, Dantas G (2014). Bacterial phylogeny structures soil resistomes across habitats. Nature 10.1038/nature13377

Freter R, Brickner H, Temme S (1986). An understanding of colonization resistance of the mammalian large intestine requires mathematical analysis. Microecology and Therapy 16: 147–155

Gao P, Munir M, Xagoraraki I (2012). Correlation of tetracycline and sulfonamide antibiotics with corresponding resistance genes and resistant bacteria in a conventional municipal wastewater treatment plant. Science of The Total Environment 421–422: 173–183 10.1016/j.scitotenv.2012.01.061

Gibson C, Kraemer SA, Klimova N, Guo B, Frigon D (2023). Antibiotic resistance gene sequencing is necessary to reveal the complex dynamics of immigration from sewers to activated sludge. Frontiers in Microbiology 14: 10.3389/fmicb.2023.1155956

Godon J-J, Arcemisbéhère L, Escudié R, Harmand J, Miambi E, Steyer J-P (2013). Overview of the oldest existing set of substrate-optimized anaerobic processes: digestive tracts. BioEnergy Research 6: 1063–1081 10.1007/s12155-013-9339-y

Goldfarb T, Kodali VK, Pujar S, Brover V, Robbertse B, Farrell CM, Oh D-H, Astashyn A, Ermolaeva O, Haddad D, Hlavina W, Hoffman J, Jackson JD, Joardar VS, Kristensen D, Masterson P, McGarvey KM, McVeigh R, Mozes E, Murphy MR, Schafer SS, Souvorov A, Spurrier B, Strope PK, Sun H, Vatsan AR, Wallin C, Webb D, Brister JR, Hatcher E, Kimchi A, Klimke W, Marchler-Bauer A, Pruitt KD, Thibaud-Nissen F, Murphy TD (2025). NCBI RefSeq: reference sequence standards through 25 years of curation and annotation. Nucleic Acids Research 53: D243–D257 10.1093/nar/gkae1038

Goulas A, Belhadi D, Descamps A, Andremont A, Benoit P, Courtois S, Dagot C, Grall N, Makowski D, Nazaret S, Nélieu S, Patureau D, Petit F, Roose-Amsaleg C, Vittecoq M, Livoreil B, Laouénan C (2020). How effective are strategies to control the dissemination of antibiotic resistance in the environment? A systematic review. Environmental Evidence 9: 4 10.1186/s13750-020-0187-x

Gullberg E, Albrecht LM, Karlsson C, Sandegren L, Andersson DI (2014). Selection of a Multidrug Resistance Plasmid by Sublethal Levels of Antibiotics and Heavy Metals. mBio 5: e01918–14 10.1128/mBio.01918-14

Gullberg E, Cao S, Berg OG, Ilbäck C, Sandegren L, Hughes D, Andersson DI (2011). Selection of Resistant Bacteria at Very Low Antibiotic Concentrations. PLOS Pathogens 7: e1002158 10.1371/journal.ppat.1002158

Guo X, Chen X, Sidikjan N, Sha R (2024). Silver nanoparticles regulate antibiotic resistance genes by shifting bacterial community and generating anti-silver genes in estuarine biofilms. Aquatic Toxicology 276: 107131 10.1016/j.aquatox.2024.107131

Herbert D, Elsworth R, Telling RC (1956). The continuous culture of bacteria; a theoretical and experimental study. J Gen Microbiol 14: 601–622

Honda R, Matsuura N, Sorn S, Asakura S, Morinaga Y, Van Huy T, Sabar MA, Masakke Y, Hara-Yamamura H, Watanabe T (2023). Transition of antimicrobial resistome in wastewater treatment plants: impact of process configuration, geographical location and season. npj Clean Water 6: 1–12 10.1038/s41545-023-00261-x

Huerta-Cepas J, Serra F, Bork P (2016). ETE 3: Reconstruction, Analysis, and Visualization of Phylogenomic Data. Molecular Biology and Evolution 33: 1635–1638 10.1093/molbev/msw046

Hyatt D, Chen G-L, LoCascio PF, Land ML, Larimer FW, Hauser LJ (2010). Prodigal: prokaryotic gene recognition and translation initiation site identification. BMC Bioinformatics 11: 119 10.1186/1471-2105-11-119

Jannasch HW (1969). Estimations of bacterial growth rates in natural waters. Journal of Bacteriology 99: 156–160

Jannasch HW, Egli T (1993). Microbial growth kinetics: a historical perspective. Antonie van Leeuwenhoek 63: 213–224 10.1007/BF00871219

Jia S, Zhang X-X, Miao Y, Zhao Y, Ye L, Li B, Zhang T (2017). Fate of antibiotic resistance genes and their associations with bacterial community in livestock breeding wastewater and its receiving river water. Water Research 124: 259–268 10.1016/j.watres.2017.07.061

Ju F, Beck K, Yin X, Maccagnan A, McArdell CS, Singer HP, Johnson DR, Zhang T, Bürgmann H (2019). Wastewater treatment plant resistomes are shaped by bacterial composition, genetic exchange, and upregulated expression in the effluent microbiomes. The ISME Journal 13: 346–360 10.1038/s41396-018-0277-8

Keskey R, Bluiminck S, Sangwan N, Meltzer R, Lam A, Thewissen R, Zaborin A, van Goor H, Zaborina O, Alverdy J (2025). Dietary impact on the gut resistome: western diet independently increases the prevalence of antibiotic resistance genes within the gut microbiota. Microbiology Spectrum 0: e02766–24 10.1128/spectrum.02766-24

Kim S, Park H, Chandran K (2010). Propensity of activated sludge to amplify or attenuate tetracycline resistance genes and tetracycline resistant bacteria: A mathematical modeling approach. Chemosphere 78: 1071–1077 10.1016/j.chemosphere.2009.12.068

Klümper U, Gionchetta G, Catão E, Bellanger X, Dielacher I, Elena AX, Fang P, Galazka S, Goryluk-Salmonowicz A, Kneis D, Okoroafor U, Radu E, Szadziul M, Szekeres E, Teban-Man A, Coman C, Kreuzinger N, Popowska M, Vierheilig J, Walsh F, Woegerbauer M, Bürgmann H, Merlin C, Berendonk TU (2024). Environmental microbiome diversity and stability is a barrier to antimicrobial resistance gene accumulation. Communications Biology 7: 1–13 10.1038/s42003-024-06338-8

Klümper U, Recker M, Zhang L, Yin X, Zhang T, Buckling A, Gaze WH (2019). Selection for antimicrobial resistance is reduced when embedded in a natural microbial community. The ISME Journal 13: 2927–2937 10.1038/s41396-019-0483-z

Knapp CW, Engemann CA, Hanson ML, Keen PL, Hall KJ, Graham DW (2008). Indirect Evidence of Transposon-Mediated Selection of Antibiotic Resistance Genes in Aquatic Systems at Low-Level Oxytetracycline Exposures. Environmental Science & Technology 42: 5348–5353 10.1021/es703199g

Kneis D, Barron M de la C, Konyali D, Westphal V, Schröder P, Westphal-Settele K, Schönfeld J, Jungmann D, Berendonk TU, Klümper U (2025). Suppressing selection for antibiotic resistance in the environment: A transparent, ecology-based approach to predicted no-effect concentrations. 2025.04.04.647007 10.1101/2025.04.04.647007

Koch AL (2001). Oligotrophs versus copiotrophs. BioEssays 23: 657–661 10.1002/bies.1091

Kurasam J, Mandal PK, Sarkar S (2022). Selective Proliferation of Antibiotic-Resistant Bacteria in the Biological Treatment Process at a Municipal Wastewater Treatment Plant in India. Journal of Environmental Engineering 148: 04022007 10.1061/(ASCE)EE.1943-7870.0001980

Kuznetsov SI, Dubinina GA, Lapteva NA (1979). Biology of oligotrophic bacteria. Annual Review of Microbiology 33: 377–387 10.1146/annurev.mi.33.100179.002113

Larsson DGJ, Gaze WH, Laxminarayan R, Topp E (2023). AMR, One Health and the environment. Nature Microbiology 1–2 10.1038/s41564-023-01351-9

Lauro FM, McDougald D, Thomas T, Williams TJ, Egan S, Rice S, DeMaere MZ, Ting L, Ertan H, Johnson J, Ferriera S, Lapidus A, Anderson I, Kyrpides N, Munk AC, Detter C, Han CS, Brown MV, Robb FT, Kjelleberg S, Cavicchioli R (2009). The genomic basis of trophic strategy in marine bacteria. Proceedings of the National Academy of Sciences of the United States of America 106: 15527–15533 10.1073/pnas.0903507106

Lawrence M, Homer-Dixon T, Janzwood S, Rockstöm J, Renn O, Donges JF (2024). Global polycrisis: the causal mechanisms of crisis entanglement. Global Sustainability 7: e6 10.1017/sus.2024.1

Lee K, Raguideau S, Sirén K, Asnicar F, Cumbo F, Hildebrand F, Segata N, Cha C-J, Quince C (2023). Population-level impacts of antibiotic usage on the human gut microbiome. Nature Communications 14: 1191 10.1038/s41467-023-36633-7

Lendenmann U, Egli T (1998). Kinetic models for the growth of Escherichia coli with mixtures of sugars under carbon-limited conditions. Biotechnology and Bioengineering 59: 99–107 10.1002/(sici)1097-0290(19980705)59:1%253C99::aid-bit13%253E3.0.co;2-y

Lessard EJ, Sieburth JMcN (1983). Survival of Natural Sewage Populations of Enteric Bacteria in Diffusion and Batch Chambers in the Marine Environment. Applied and Environmental Microbiology 45: 950–959

Letten AD, Hall AR, Levine JM (2021). Using ecological coexistence theory to understand antibiotic resistance and microbial competition. Nature Ecology & Evolution 5: 431–441 10.1038/s41559-020-01385-w

Li J, Mau RL, Dijkstra P, Koch BJ, Schwartz E, Liu X-JA, Morrissey EM, Blazewicz SJ, Pett-Ridge J, Stone BW, Hayer M, Hungate BA (2019). Predictive genomic traits for bacterial growth in culture versus actual growth in soil. The ISME Journal 13: 2162–2172 10.1038/s41396-019-0422-z

Li S, Zhou X, Liu L, Su Z, Zhao J, Zhang J, Cai Z, Peñuelas J, Huang X (2025). Plant Diversity Reduces the Risk of Antibiotic Resistance Genes in Agroecosystems. Advanced Science n/a: 2410990 10.1002/advs.202410990

Lin Q, Xavier BB, Alako BTF, Mitchell AL, Rajakani SG, Glupczynski Y, Finn RD, Cochrane G, Malhotra-Kumar S (2022). Screening of global microbiomes implies ecological boundaries impacting the distribution and dissemination of clinically relevant antimicrobial resistance genes. Communications Biology 5: 1–9 10.1038/s42003-022-04187-x

Lundström SV, Östman M, Bengtsson-Palme J, Rutgersson C, Thoudal M, Sircar T, Blanck H, Eriksson KM, Tysklind M, Flach C-F, Larsson DGJ (2016). Minimal selective concentrations of tetracycline in complex aquatic bacterial biofilms. Science of The Total Environment 553: 587–595 10.1016/j.scitotenv.2016.02.103

Luo T, Dai X, Chen Z, Wu L, Wei W, Xu Q, Ni B-J (2022). Different microplastics distinctively enriched the antibiotic resistance genes in anaerobic sludge digestion through shifting specific hosts and promoting horizontal gene flow. Water Research 228: 119356 10.1016/j.watres.2022.119356

Ma Y, Qiao Y, Zhang X, Ye L (2024). Filamentous bacteria-induced sludge bulking can alter antibiotic resistance gene profiles and increase potential risks in wastewater treatment systems. Environment International 190: 108920 10.1016/j.envint.2024.108920

Mao D, Yu S, Rysz M, Luo Y, Yang F, Li F, Hou J, Mu Q, Alvarez PJJ (2015). Prevalence and proliferation of antibiotic resistance genes in two municipal wastewater treatment plants. Water Research 85: 458–466 10.1016/j.watres.2015.09.010

Marsh PD (1995). The role of continuous culture in modeling the human microflora. Journal of Chemical Technology and Biotechnology 64: 1–9

Martiny H-M, Munk P, Brinch C, Aarestrup FM, Petersen TN (2022). A curated data resource of 214K metagenomes for characterization of the global antimicrobial resistome. PLOS Biology 20: e3001792 10.1371/journal.pbio.3001792

McArthur AG, Waglechner N, Nizam F, Yan A, Azad MA, Baylay AJ, Bhullar K, Canova MJ, De Pascale G, Ejim L, Kalan L, King AM, Koteva K, Morar M, Mulvey MR, O’Brien JS, Pawlowski AC, Piddock LJV, Spanogiannopoulos P, Sutherland AD, Tang I, Taylor PL, Thaker M, Wang W, Yan M, Yu T, Wright GD (2013). The Comprehensive Antibiotic Resistance Database. Antimicrobial Agents and Chemotherapy 57: 3348–3357 10.1128/aac.00419-13

Muñoz-Aguayo J, Lang KS, LaPara TM, González G, Singer RS (2007). Evaluating the Effects of Chlortetracycline on the Proliferation of Antibiotic-Resistant Bacteria in a Simulated River Water Ecosystem. Applied and Environmental Microbiology 73: 5421–5425 10.1128/AEM.00708-07

Murray AK, Stanton IC, Wright J, Zhang L, Snape J, Gaze WH (2020). The ‘SELection End points in Communities of bacTeria’ (SELECT) Method: A Novel Experimental Assay to Facilitate Risk Assessment of Selection for Antimicrobial Resistance in the Environment. Environmental Health Perspectives 128: 107007 10.1289/EHP6635

Murray AK, Zhang L, Yin X, Zhang T, Buckling A, Snape J, Gaze WH (2018). Novel Insights into Selection for Antibiotic Resistance in Complex Microbial Communities. mBio 9: 10.1128/mBio.00969-18

Murray CJ, Ikuta KS, Sharara F, Swetschinski L, Aguilar GR, Gray A, et al. (2022). Global burden of bacterial antimicrobial resistance in 2019: a systematic analysis. The Lancet 399: 629–655 10.1016/S0140-6736(21)02724-0

Naghavi M, Vollset SE, Ikuta KS, Swetschinski LR, Gray AP, Wool EE, et al. (2024). Global burden of bacterial antimicrobial resistance 1990–2021: a systematic analysis with forecasts to 2050. The Lancet 404: 1199–1226 10.1016/S0140-6736(24)01867-1

Nair RR, Andersson DI (2023). Interspecies interaction reduces selection for antibiotic resistance in Escherichia coli. Communications Biology 6: 1–9 10.1038/s42003-023-04716-2

Ondov BD, Bergman NH, Phillippy AM (2011). Interactive metagenomic visualization in a Web browser. BMC Bioinformatics 12: 385 10.1186/1471-2105-12-385

Ott A, Quintela-Baluja M, Zealand AM, O’Donnell G, Haniffah MRM, Graham DW (2021). Improved quantitative microbiome profiling for environmental antibiotic resistance surveillance. Environmental Microbiome 16: 21 10.1186/s40793-021-00391-0

Ovis-Sánchez JO, Vital-Jácome M, Buitrón G, Cervantes-Avilés P, Carrillo-Reyes J (2025). Antibiotic resistance reduction mechanisms during thermophilic anaerobic digestion of microalgae-bacteria aggregates. Bioresource Technology 419: 132037 10.1016/j.biortech.2025.132037

Pallares-Vega R, Blaak H, van der Plaats R, de Roda Husman AM, Hernandez Leal L, van Loosdrecht MCM, Weissbrodt DG, Schmitt H (2019). Determinants of presence and removal of antibiotic resistance genes during WWTP treatment: A cross-sectional study. Water Research 161: 319–328 10.1016/j.watres.2019.05.100

Park J-H, Bae K-S, Kang J, Park E-R, Yoon J-K (2024). Comprehensive Study of Antibiotic Resistance in Enterococcus spp.: Comparison of Influents and Effluents of Wastewater Treatment Plants. Antibiotics 13: 1072 10.3390/antibiotics13111072

Poindexter JS (1981). Oligotrophy - Fast and Famine Existence. In: Alexander M (ed), Advances in Microbial Ecology. Springer US, Boston, MA, pp. 63–89

Props R, Kerckhof F-M, Rubbens P, De Vrieze J, Hernandez Sanabria E, Waegeman W, Monsieurs P, Hammes F, Boon N (2017). Absolute quantification of microbial taxon abundances. The ISME Journal 11: 584–587 10.1038/ismej.2016.117

Quinlan EL, Nietch CT, Blocksom K, Lazorchak JM, Batt AL, Griffiths R, Klemm DJ (2011). Temporal Dynamics of Periphyton Exposed to Tetracycline in Stream Mesocosms. Environmental Science & Technology 45: 10684–10690 10.1021/es202004k

Quintela-Baluja M, Frigon D, Abouelnaga M, Jobling K, Romalde JL, Gomez Lopez M, Graham DW (2021). Dynamics of integron structures across a wastewater network – Implications to resistance gene transfer. Water Research 206: 117720 10.1016/j.watres.2021.117720

Roller BRK, Stoddard SF, Schmidt TM (2016). Exploiting rRNA operon copy number to investigate bacterial reproductive strategies. Nature Microbiology 1: 1–7 10.1038/nmicrobiol.2016.160

Schut F, Prins RA, Gottschal JC (1997). Oligotrophy and pelagic marine bacteria: Facts and fiction. Aquatic Microbial Ecology 12: 177–202

Seemann T (2018). Barrnap: BAsic Rapid Ribosomal RNA Predictor.

Segura Munoz RR, Mantz S, Martínez I, Li F, Schmaltz RJ, Pudlo NA, Urs K, Martens EC, Walter J, Ramer-Tait AE (2022). Experimental evaluation of ecological principles to understand and modulate the outcome of bacterial strain competition in gut microbiomes. The ISME Journal 16: 1594–1604 10.1038/s41396-022-01208-9

Soler-Bistué A, Couso LL, Sánchez IE (2023). The evolving copiotrophic/oligotrophic dichotomy: From Winogradsky to physiology and genomics. Environmental Microbiology 25: 1232–1237 10.1111/1462-2920.16360

Stanton IC, Murray AK, Zhang L, Snape J, Gaze WH (2020). Evolution of antibiotic resistance at low antibiotic concentrations including selection below the minimal selective concentration. Communications Biology 3: 1–11 10.1038/s42003-020-01176-w

Tian Z, Palomo A, Zhang H, Luan X, Liu R, Awad M, Smets BF, Zhang Y, Yang M (2020). Minimum influent concentrations of oxytetracycline, streptomycin and spiramycin in selecting antibiotic resistance in biofilm type wastewater treatment systems. Science of The Total Environment 720: 137531 10.1016/j.scitotenv.2020.137531

Tilman D (1982). Resource Competition and Community Structure Princeton University Press, New Jersey

United Nations Environment Programme (UNEP) (2023). Bracing for Superbugs: Strengthening environmental action in the One Health response to antimicrobial resistance Geneva

Vandeputte D, Kathagen G, D’hoe K, Vieira-Silva S, Valles-Colomer M, Sabino J, Wang J, Tito RY, De Commer L, Darzi Y, Vermeire S, Falony G, Raes J (2017). Quantitative microbiome profiling links gut community variation to microbial load. Nature 551: 507–511 10.1038/nature24460

Wimpenny JWT (1988). CRC Handbook of Laboratory Model Systems for Microbial Ecosystems CRC Press, Boca Raton, FL, USA

Woodworth MH, Conrad RE, Haldopoulos M, Pouch SM, Babiker A, Mehta AK, Sitchenko KL, Wang CH, Strudwick A, Ingersoll JM, Philippe C, Lohsen S, Kocaman K, Lindner BG, Hatt JK, Jones RM, Miller C, Neish AS, Friedman-Moraco R, Karadkhele G, Liu KH, Jones DP, Mehta CC, Ziegler TR, Weiss DS, Larsen CP, Konstantinidis KT, Kraft CS (2023). Fecal microbiota transplantation promotes reduction of antimicrobial resistance by strain replacement. Science Translational Medicine 15: eabo2750 10.1126/scitranslmed.abo2750

World Economic Forum (WEF) (2023). Global Risks Report 2023 World Economic Forum, Geneva

Wu D, Zhao J, Su Y, Yang M, Dolfing J, Graham DW, Yang K, Xie B (2023). Explaining the resistomes in a megacity’s water supply catchment: Roles of microbial assembly-dominant taxa, niched environments and pathogenic bacteria. Water Research 228: 119359 10.1016/j.watres.2022.119359

Zhao Y, Su J-Q, An X-L, Huang F-Y, Rensing C, Brandt KK, Zhu Y-G (2018). Feed additives shift gut microbiota and enrich antibiotic resistance in swine gut. Science of The Total Environment 621: 1224–1232 10.1016/j.scitotenv.2017.10.106

Zhou Z-C, Zheng J, Wei Y-Y, Chen T, Dahlgren RA, Shang X, Chen H (2017). Antibiotic resistance genes in an urban river as impacted by bacterial community and physicochemical parameters. Environmental Science and Pollution Research 24: 23753–23762 10.1007/s11356-017-0032-0

Zhu C, Wu L, Ning D, Tian R, Gao S, Zhang B, Zhao J, Zhang Ya, Xiao N, Wang Y, Brown MR, Tu Q, Ju F, Wells GF, Guo J, He Z, Nielsen PH, Wang A, Zhang Yu, Chen T, He Q, Criddle CS, Wagner M, Tiedje JM, Curtis TP, Wen X, Yang Y, Alvarez-Cohen L, Stahl DA, Alvarez PJJ, Rittmann BE, Zhou J (2025). Global diversity and distribution of antibiotic resistance genes in human wastewater treatment systems. Nature Communications 16: 4006 10.1038/s41467-025-59019-3

