## Supplementary_material for "Inferring antibiotic resistance selection in the environment can be confounded by correlations between resistance genes and unrelated functional traits"

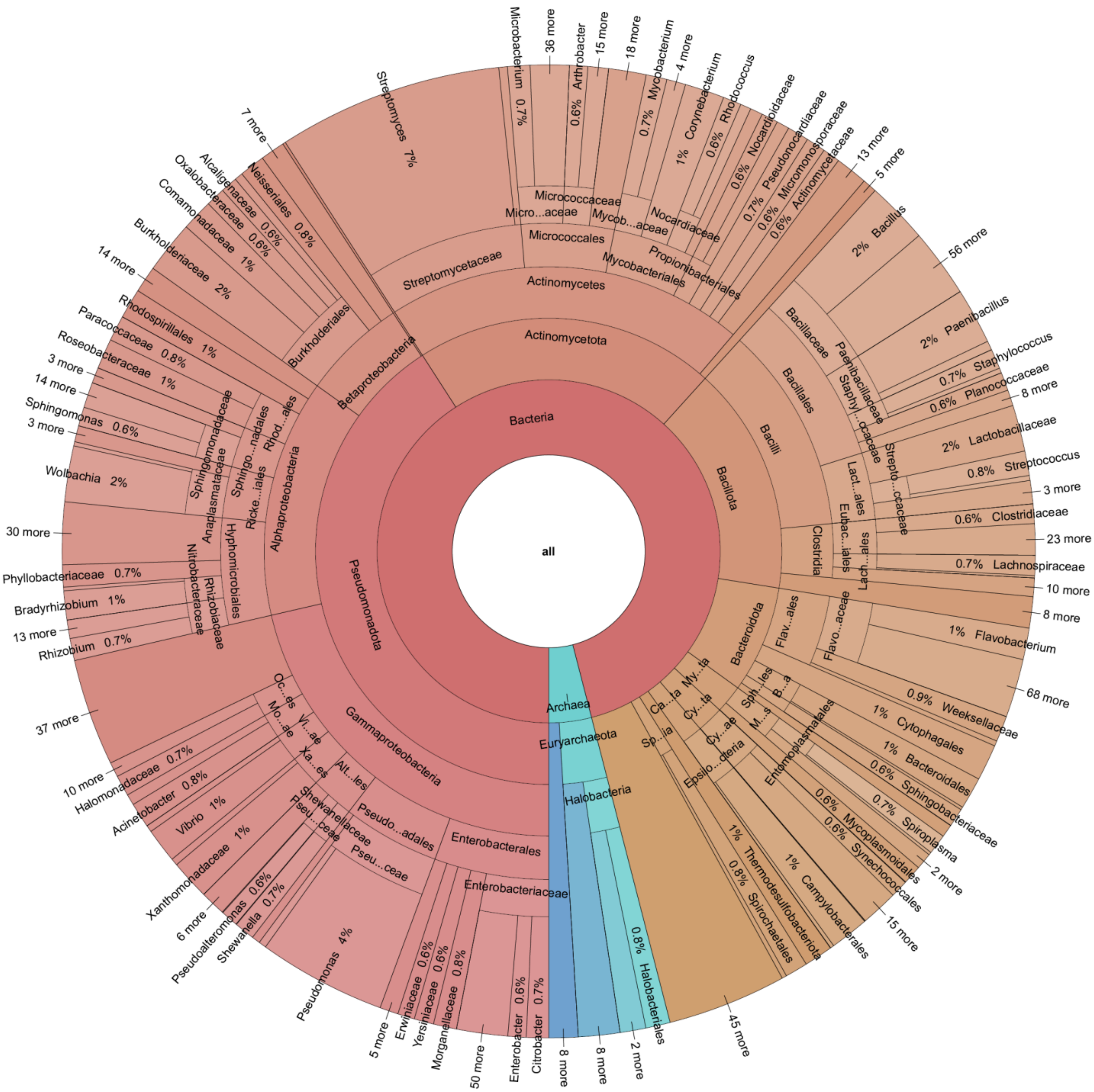


**Figure S1: The RefSeq database is biased towards easily cultured, fast growing bacteria, in particular pathogens.** The Krona figure shows that few phyla contain the majority of known genomes (Pseudomonadota, Actinomycetota and Bacillota) in contrast to many phyla with few known genomes.

**Table S1:** Parameters for the model for both cases. For equations, see the next section.

| Parameter (units) | Symbol | Case E2L (Switch from Environment (chemostat) to Laboratory (batch culture) | Case L2H (Switch from chemostat at Low dilution rate to High dilution rate) |
|---|---|---|---|
| **Maximum growth rate (per min)** | $\mu_{max,i}$ | [0.016, 0.015, 0.014, 0.00103, 0.00102, 0.001] | [0.0102, 0.0101, 0.0100, 0.001062, 0.00103, 0.001] |
| **Affinity matrix L/(mg·min)** Affinity is the ratio $\mu_{max}/K_s$, i.e, the slope of the curve at the origin | $a_{ij}$ | [0.0104 0.01 0.01 0.0105 0.01 0.01<br>0.01 0.0106 0.01 0.01 0.0107 0.01<br>0.01 0.01 0.0108 0.01 0.01 0.0109<br>0.101 0.1 0.1 0.102 0.1 0.1<br>0.1 0.103 0.1 0.1 0.104 0.1<br>0.1 0.1 0.105 0.1 0.1 0.106] | [0.010305 0.01 0.01 0.0104 0.01 0.01<br>0.01 0.0104 0.01 0.01 0.0105 0.01<br>0.01 0.01 0.0105 0.01 0.01 0.0106<br>0.101 0.1 0.1 0.102 0.1 0.1<br>0.1 0.1038 0.1 0.1 0.105 0.1<br>0.1 0.1 0.1068 0.1 0.1 0.1079] |
| **Yield coefficient** | $Y_{ij}$ | 0.5 (constant for all bacteria and substrates) | 0.5 (constant for all bacteria and substrates) |
| **Dilution rate (chemostat) (per min)** | $D$ | Initially: min $(\mu_{max,i})/4$ = 0.00025<br>After switch: 0 | Initially: min $(\mu_{max,i})/4$ = 0.00025<br>After switch: 10 x initial D |
| **Immigration rate (mg/(L min))** | $r$ | 0.01 × max$(\mu_{max,i})$ = 0.00016 | 0.01 × max$(\mu_{max,i})$ |
| **Reservoir substrate concentration (mg/L)** | $s_{o,j}$ | 1,000 for all substrates | 1,000 for all substrates |
| **Batch substrate addition (mg/L)** | $s_b$ | Injected once at the time of switching: 1000 | N/A |
| **Initial biomass concentration (mg/L)** | $x_{i,0}$ | 50 for all bacterial species | 50 for all bacterial species |
| **Initial substrate concentration (mg/L)** | $s_{j,0}$ | Equal to $s_0$ | Equal to $s_0$ |
| **Simulation time** | $T$ | 10 × (1/D) = 40,000 min | 16,000 × $(1/D)$ |
| **Switch time (to batch/different dilution rate)** | $t_{switch}$ | $T$ × (3/4) = 30,000 min | $T/2$ |

### System of ODEs

This section provides the system of ordinary differential equations (ODEs) for studying the two cases, the shift in bacterial communities from a chemostat at low dilution rate to higher dilution rate or the switch from chemostat to a batch reactor environment. The equations for the chemostat contain the batch reactor as a special case with dilution rate set to zero.

Equations are general, but the parameters in **Table S1** are for *n* = 6 bacterial species (i=1, 2, …, 6) and *k* = 6 substrates (j=1, 2, …, 6).

The specific growth rate (based on an adaptation of Monod kinetics to growth on multiple substitutable substrates by Lendenmann & Egli (1998), their Eq. 17) for bacterial species *i* is given by:

$$\mu_i = \frac{\mu_{max,i}\ \sum_{j=1}^{k} a_{ij} s_j}{\mu_{max,i}\ +\ \sum_{j=1}^{k} a_{ij} s_j}$$

Bacterial biomass (*x*) dynamics (D is set to 0 for batch cultures):

$$\frac{dx_i}{dt} = (\mu_i - D)\ x_i + r$$

Substrate (*s*) dynamics (D is set to 0 for batch cultures):

$$\frac{ds_j}{dt} = D(s_{o,j} - s_j) - \sum_{i=1}^{n} Y_{ij}\ \mu_i\ x_i$$

### Notes

Immigration Rate: The immigration rate term *r* is added to the biomass equations but does not affect substrates directly. Immigration happens constantly for both case studies before and after the shift.

Resource competition: Substrate consumption depends on $Y_{ij}$ , which is constant here for simplicity.

For the L2H case (switch from chemostat at Low dilution rate to High dilution rate):

1. Switch in dilution rate: At t = t_switch, D switches to 10 x Original D
2. Affinity Matrix: Adjustments in $a_{ij}$ for higher dilution rates (slightly higher affinities in some cases).

For the E2L case (switch from Environment (chemostat) to Laboratory (batch culture):

1. Switch in dynamics: At t = t_switch, the system switches from a chemostat (constant dilution rate D > 0) to a batch reactor (D = 0).
2. At the same time, a large amount of all six substrates is given once only (substrate concentrations are set to the reservoir concentrations of 1,000 mg/L).

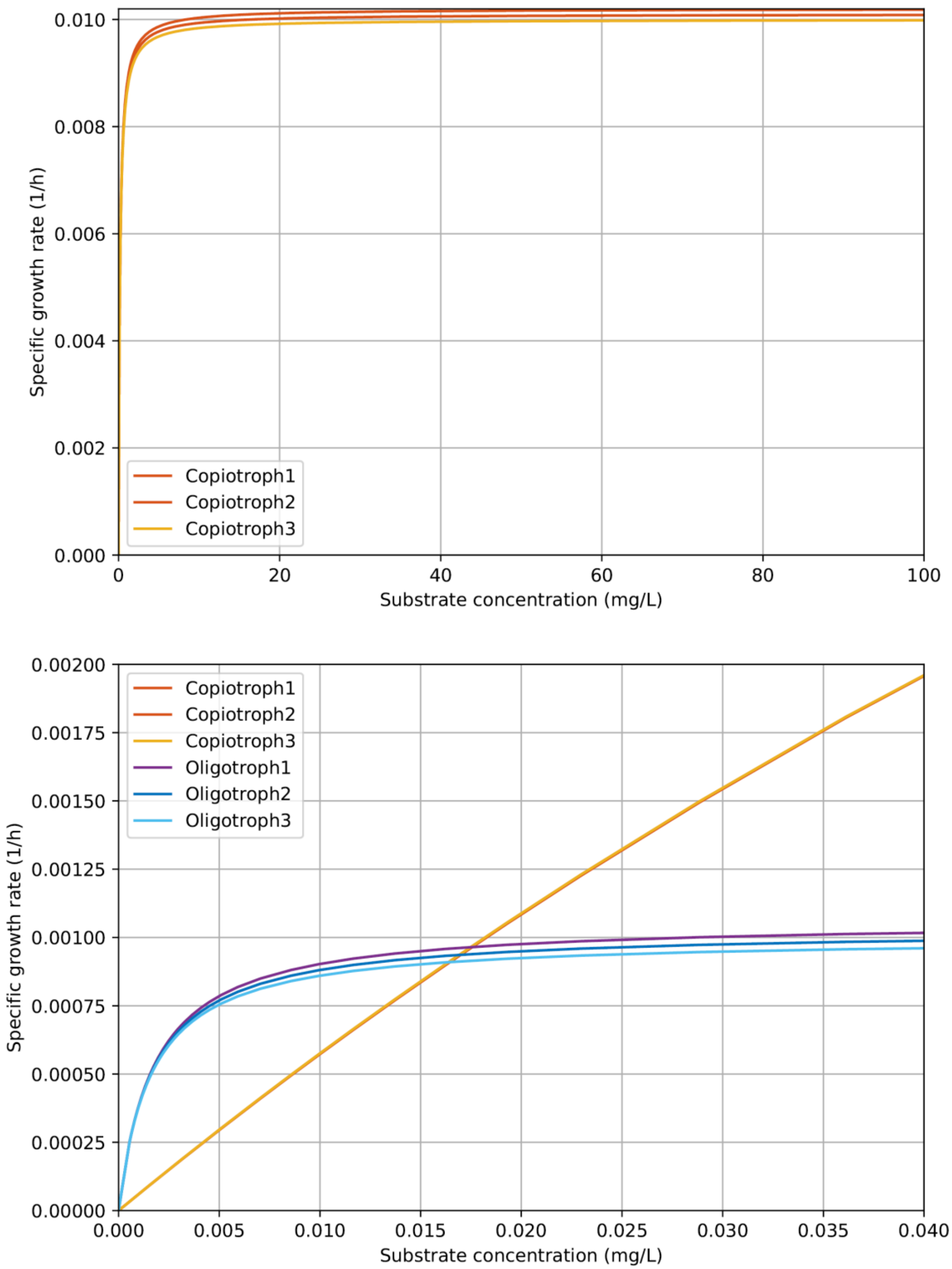


**Figure S2: Growth kinetics for the low to high dilution rate shift (L2H) case study.** The kinetics of the three copiotrophic species were similar but not identical, with much higher maximum specific growth rates and lower substrate affinities than the three oligotrophic species, which were again similar with each other but not identical. At substrate concentrations below ~0.017 mg/L, the oligotrophs had higher specific growth rates than the copiotrophs. The top panel shows the copiotrophs and the bottom panel zoomed to show all species.

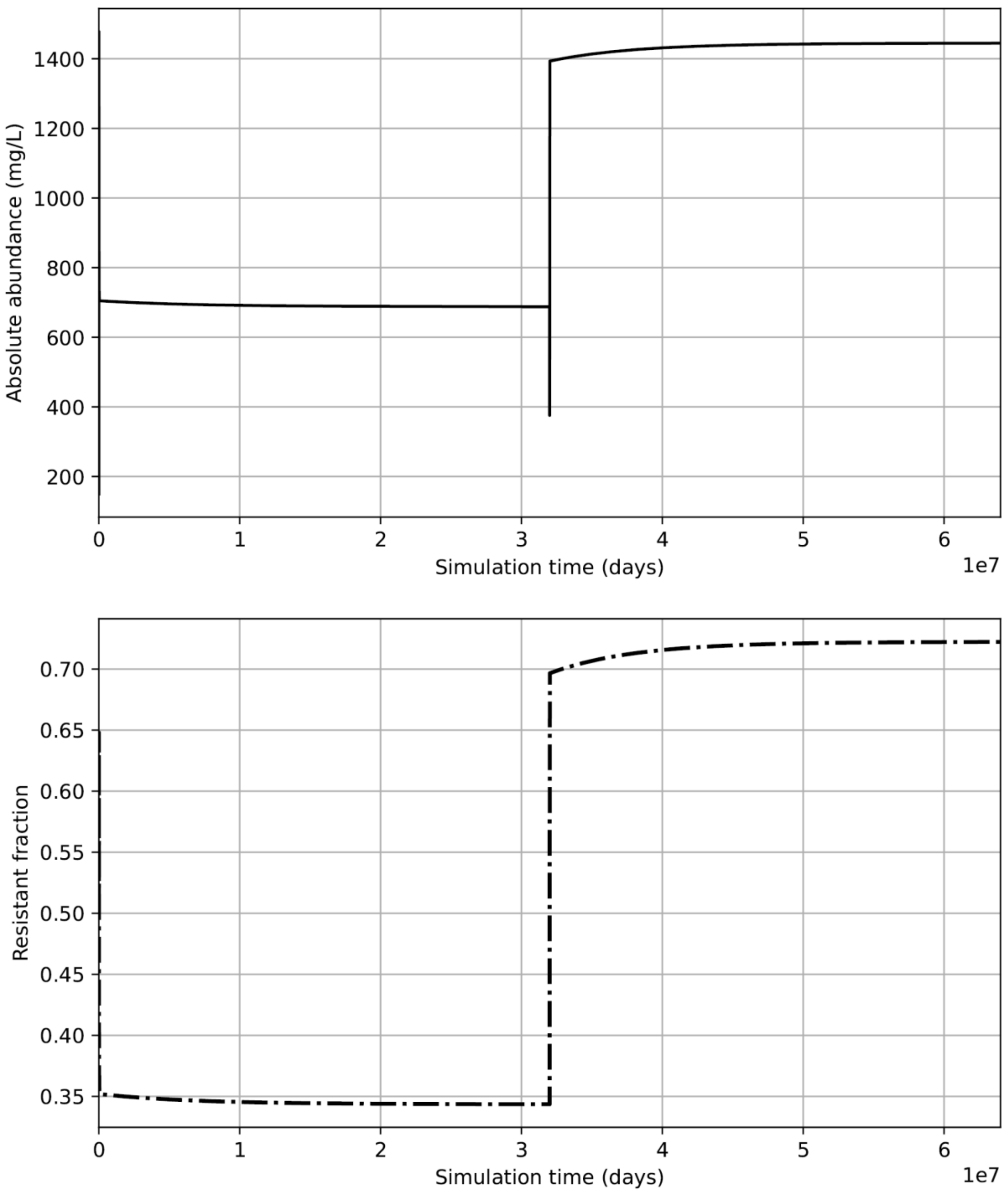


**Figure S3: Both the absolute and relative abundance of all resistant species in the community shifted up when the dilution rate and thus specific growth rate shifted up.** (Top panel) Absolute abundance of the resistance gene. (Bottom panel) Relative abundance or fraction of the resistance gene in the community. Same simulation as shown in **Figure 2**.

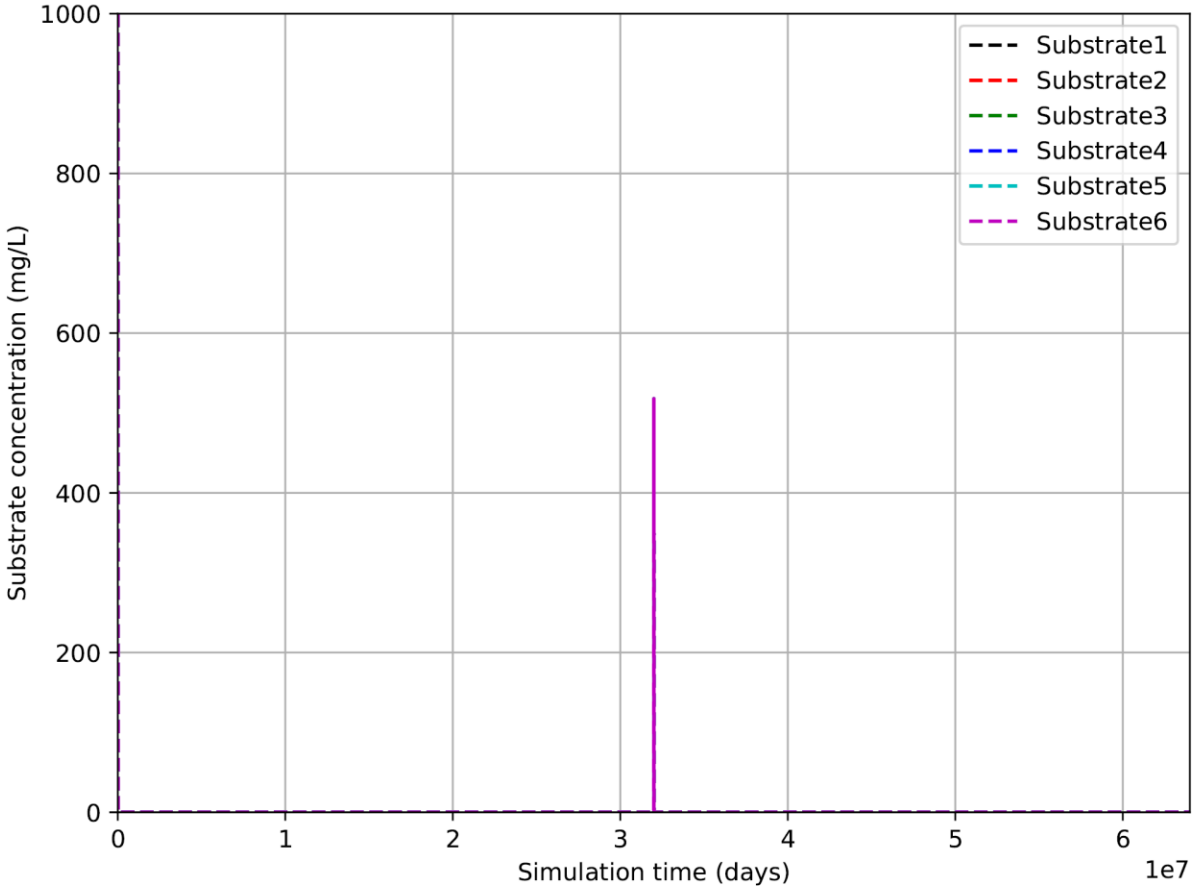


**Figure S4: Dynamics of substrate concentrations in the chemostat for the low to high dilution rate shift (L2H) case study.** Apart from the initial phase and immediately following the upshift, the six substrate concentrations remained at low levels. Same simulation as shown in **Figure 2**.

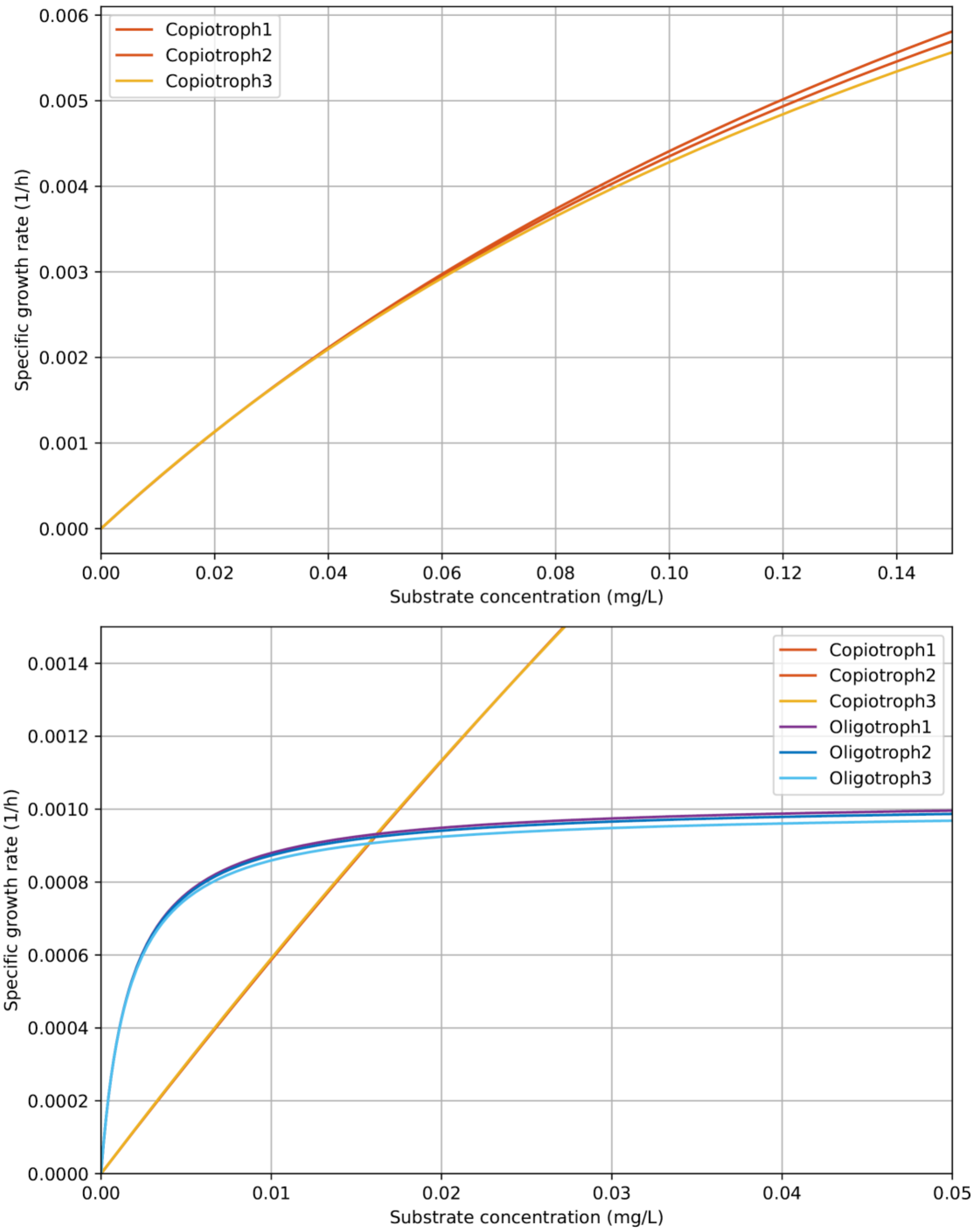


**Figure S5: Growth kinetics for the environment to laboratory shift case study (E2L).** The kinetics of the three copiotrophic species were similar but not identical, with much higher maximum specific growth rates and lower substrate affinities than the three oligotrophic species, which were again similar with each other but not identical. At substrate concentrations below ~0.017 mg/L, the oligotrophs had higher specific growth rates than the copiotrophs. The top panel shows the copiotrophs and the bottom panel zoomed to show all species. Kinetics for this E2L case study were similar to the L2H case study.

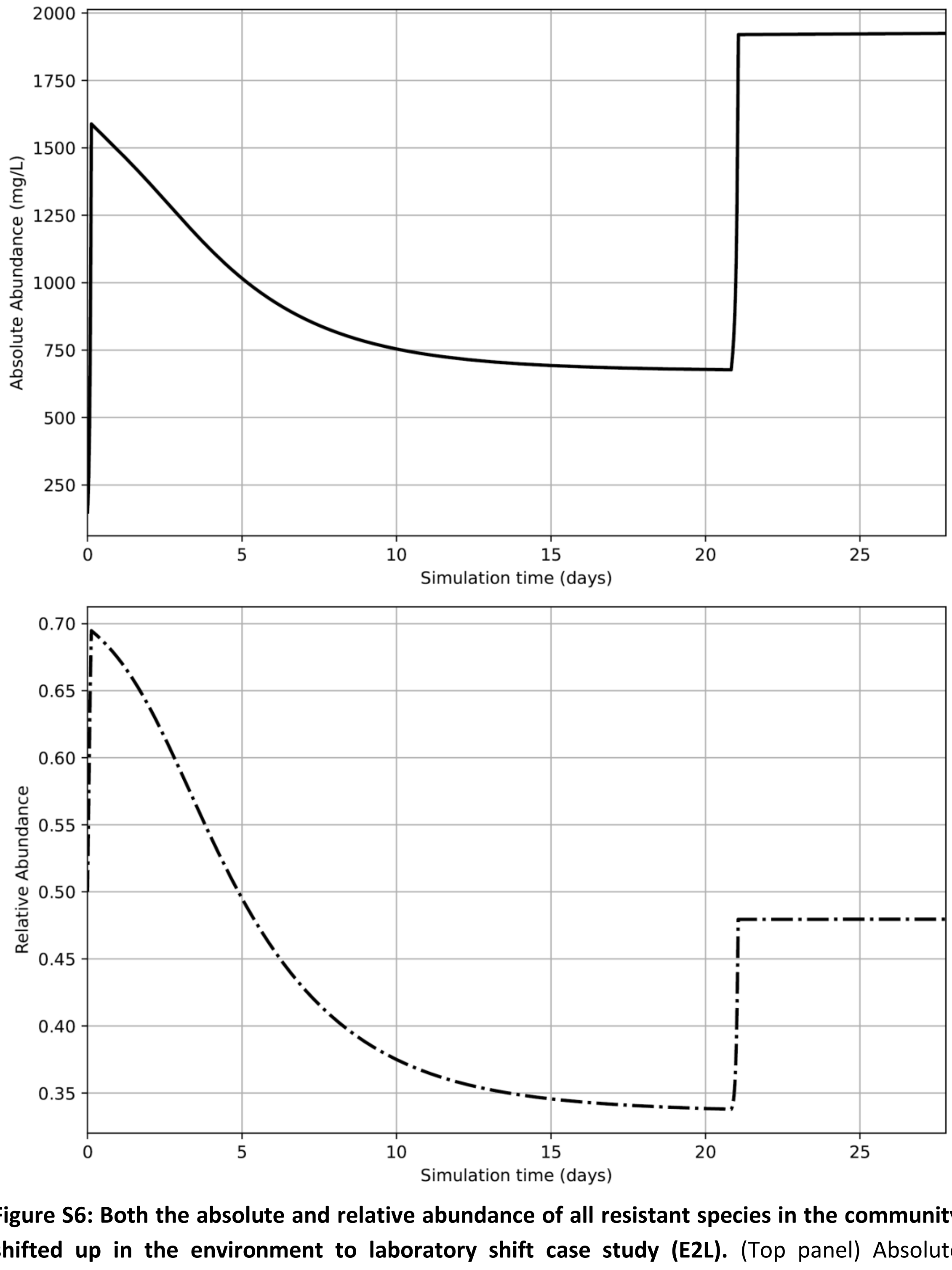


**Figure S6: Both the absolute and relative abundance of all resistant species in the community shifted up in the environment to laboratory shift case study (E2L).** (Top panel) Absolute abundance of the resistance gene. (Bottom panel) Relative abundance or fraction of the resistance gene in the community. Compared to the L2H case study, the absolute abundance increased more strongly than the relative abundance as the total population density increased. Same simulation as shown in **Figure 2**.

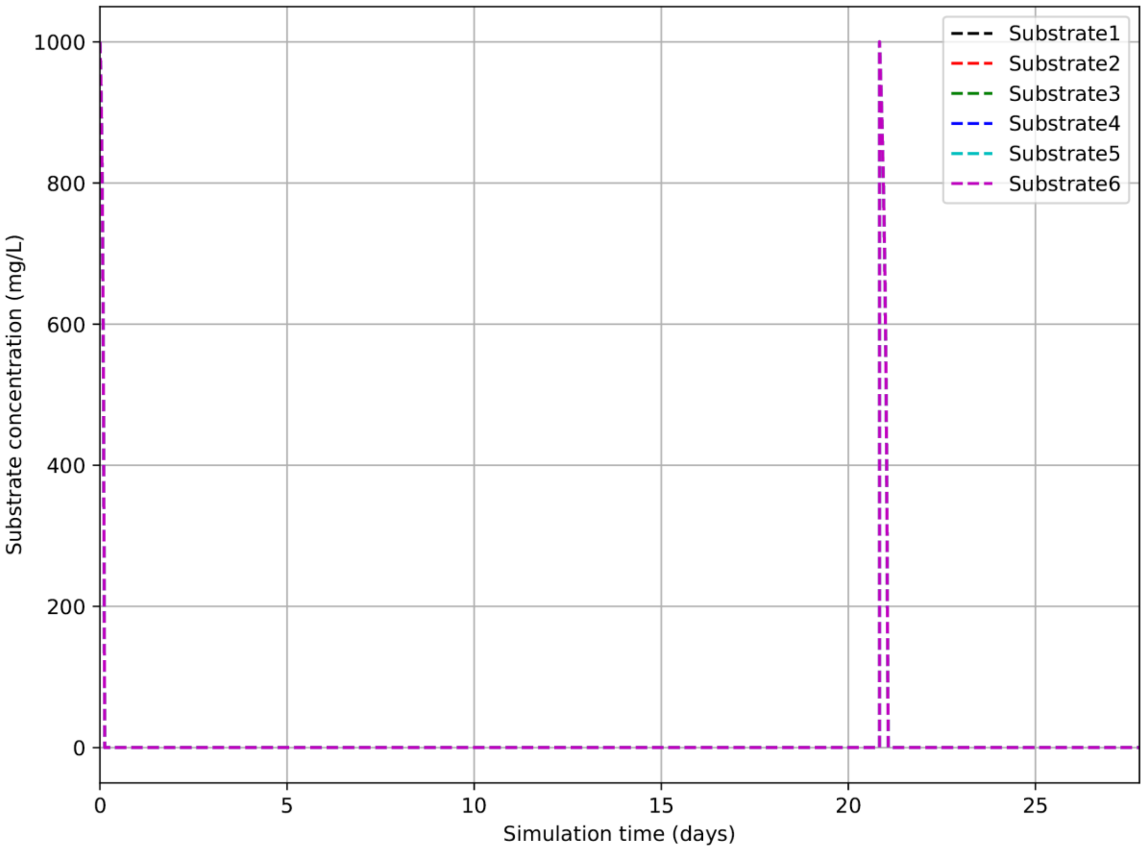


74 **Figure S7: Dynamics of substrate concentrations in the chemostat for the environment to**
75 **laboratory shift case study (E2L).** Apart from the initial phase and immediately following the
76 upshift, the six substrate concentrations remained at low levels. Same simulation as shown in
77 **Figure 3**.

### References

Lendenmann U, Egli T (1998). Kinetic models for the growth of *Escherichia coli* with mixtures of sugars under carbon-limited conditions. *Biotechnology and Bioengineering* **59**: 99–107 https://doi.org/10.1002/(sici)1097-0290(19980705)59:1<99::aid-bit13>3.0.co;2-y
